# RAS/PI3K pathway mutations sensitise epithelial ovarian cancer cells to a PARP/NAMPT inhibitor combination

**DOI:** 10.1101/2024.10.15.618473

**Authors:** Michael Gruet, Yitao Xu, Lyutong An, Yurui Ma, Cristina Balcells, Katie Tyson, Sarah Spear, Yuewei Xu, Flora McKinney, Julia Babuta, Chandler Bray, Chiharu Wickremesinghe, Alexandros P. Siskos, Anke M. Nijhuis, Edward W. Tate, Iain A. McNeish, Adrian Benito, Hector C. Keun

**Affiliations:** Department of Surgery and Cancer, Imperial College London, London W12 0NN, UK; Department of Chemistry, Molecular Sciences Research Hub, Imperial College London, London W12 0BZ, UK; Wisdom Lake Academy of Pharmacy, Xi’an Jiaotong-Liverpool University, Suzhou 215123, China; Department of Metabolism, Digestion and Reproduction, Burlington Danes, Imperial College London, London W12 OTN

**Keywords:** PARP1, NAMPT, NAD^+^, ROS, RAS, PTEN, PI3K

## Abstract

The combination of PARP and NAMPT inhibitors (PARPi/NAMPTi) has been explored for the treatment of TNBC, Ewing Sarcoma and high grade serous carcinoma (HGSC). However, dose limiting toxicity has hampered NAMPTi in clinical trials. To maximise the therapeutic window, we set out to identify predictive genomic biomarkers. Bioinformatic analysis and screening of a panel of epithelial ovarian cancer (EOC) cell lines revealed that cells with RAS/PI3K pathway mutations were sensitive to the NAMPTi FK866. Activity of olaparib and FK866 was associated with a reduction in nicotinamide mononucleotide (NMN) and the PARP substrate nicotinamide adenine dinucleotide (NAD^+^), with coincident increases in ROS production, DNA damage and apoptosis induction. Caspase 3/7 activity was upregulated to a greater extent in RAS/PI3K mutant cell lines. Finally, the combination significantly reduced omental tumour weight and increased overall survival in mice injected with ID8 *Trp53^-/-^;Pten^-/-^* cells. This study highlights the potential of the PARPi/NAMPTi combination in RAS/PI3K pathway mutant EOC.

## INTRODUCTION

Ovarian cancer is the second most commonly diagnosed gynaecological cancer in the UK^1^, and is the most common cause of gynaecological cancer deaths^2^. The most common ovarian cancer subtype, high grade serous carcinoma (HGSC), frequently harbours mutations in *BRCA1/2*, leading to a homologous recombination deficiency (HRD). HRD tumours are exquisitely sensitive to inhibitors of poly(ADP-ribose) polymerase (PARP), such as olaparib^3,4^. PARP inhibitors (PARPi) are approved in BRCA-mutated (BRCAm) patients for front-line maintenance therapy and maintenance in the recurrent setting^4^. However, evidence is emerging suggesting that patients could benefit from PARPi-therapy independent of BRCAm status^5^. Despite this, the majority of patients will relapse within 3 years while on PARPi maintenance^6^, highlighting the need to identify combination therapies to tackle treatment resistance.

PARPs regulate cellular processes, such as DNA repair, by catalysing post-translational ADP-ribosylation. This involves the covalent attachment of one or more ADP-ribose units onto targets proteins, using nicotinamide adenine dinucleotide (NAD^+^) as a substrate^7^. The majority of NAD^+^ is supplied through the NAD^+^ salvage pathway, and the rate limiting step is catalysed by the enzyme nicotinamide phosphoribosyltransferase (NAMPT). NAMPT converts nicotinamide into nicotinamide mononucleotide (NMN), which is then used to synthesise NAD^+8^. Because of the link between NAD^+^ metabolism and PARP catalytic activity, groups have investigated whether use of a NAMPT inhibitor (NAMPTi) and PARPi combination could be beneficial. Encouragingly, PARPi/NAMPTi combinations have shown synergistic activity in triple-negative breast cancer (TNBC)^9^, Ewing’s Sarcoma^10^, and HGSC^11^. However NAMPTis, such as FK866^12^, have not proved successful as monotherapies in clinical trials largely because of toxicity^13^. Considering this, there is value in identifying predictive biomarkers to improve the therapeutic window and reduce NAMPTi-associated toxicity^13^. To date only *BRCA1/2* loss has been identified as a potential biomarker for the combination^9^.

Interestingly, a post-hoc exploratory biomarker analysis from the ARIEL2 study revealed that rucaparib-treated patients harbouring alterations in genes involved in the RAS pathway or PI3K/AKT signalling (‘RAS/PI3K-mutant’) had poorer outcomes when compared to other groups^14^. RAS proteins exert their functions through several effector pathways, particularly PI3K signalling^15^. Given these pathways are frequently mutated in up to 45% of HGSC cases^16^, and in other EOC subtypes^17^, there is a need to identify therapeutic strategies that can extend the benefit of PARPis within this group of patients.

Hyperactivation of RAS/PI3K signalling results in uncontrolled proliferation and increased demands on cellular metabolism. In this context, NAD^+^ recycling must increase to sustain high proliferation rates, because NAD^+^ plays an essential role as a coenzyme in key cellular metabolic pathways, such as glycolysis and the TCA cycle^18^. Interestingly, a relationship between the NAD^+^ salvage pathway and RAS/PI3K signalling is emerging. *KRAS*-mutant tumours have been shown to possess lower intracellular NAD^+^ concentrations^19,20^, reflecting increased metabolic demand. PTEN expression inversely correlates with *NAMPT* expression^21^, and oncogenic *BRAF^V600E^* mutations can induce NAMPT overexpression, leading to NAMPTi sensitivity^22,23^. Considering this, we hypothesised that RAS/PI3K-mutant EOC cells have an increased demand for NAD^+^ that can be exploited with a NAMPTi, and consequently may exhibit greater responses to treatment with a PARPi/NAMPTi combination. To address this, we assessed the efficacy of a PARPi/NAMPTi combination *in vitro* using a panel of EOC cell lines, and *in vivo* using the ID8-*Trp53^-/-^; Pten^-/-^* model^24,25^.

Our experiments demonstrate that RAS/PI3K-mutant EOC cells are sensitive to FK866, and benefit from combined treatment with olaparib and FK866. The combination has anti-cancer activity in a murine model of HGSC with a *Pten* deletion. Together our data demonstrate the utility of a PARPi/NAMPTi combination in EOC models with genomic alterations in RAS/PI3K-pathways and suggests these mutations could be used as a predictive biomarker.

## RESULTS

### RAS/PI3K pathway mutant EOC cell lines are sensitive to the NAMPT inhibitor FK866

To investigate the role of NAMPT in EOC, data from the Genomics of Drug Sensitivity in Cancer (GDSC) resource (https://www.cancerrxgene.org/) were analysed. This analysis demonstrated that ovarian cancer cell lines are not intrinsically sensitive to the NAMPTi FK866 (**Fig. 1a**). However, when ovarian cancer cells were segregated based on their *KRAS* mutational status, FK866-sensitivity (IC_50_) was more comparable to the most sensitive cell types (highly sensitive blood cancers (**Fig. 1b**)). A significant difference in FK866-sensitivity could be observed between *KRAS*-wildtype and -mutant ovarian cancer cells (*P* = 0.001).

**Fig. 1:**
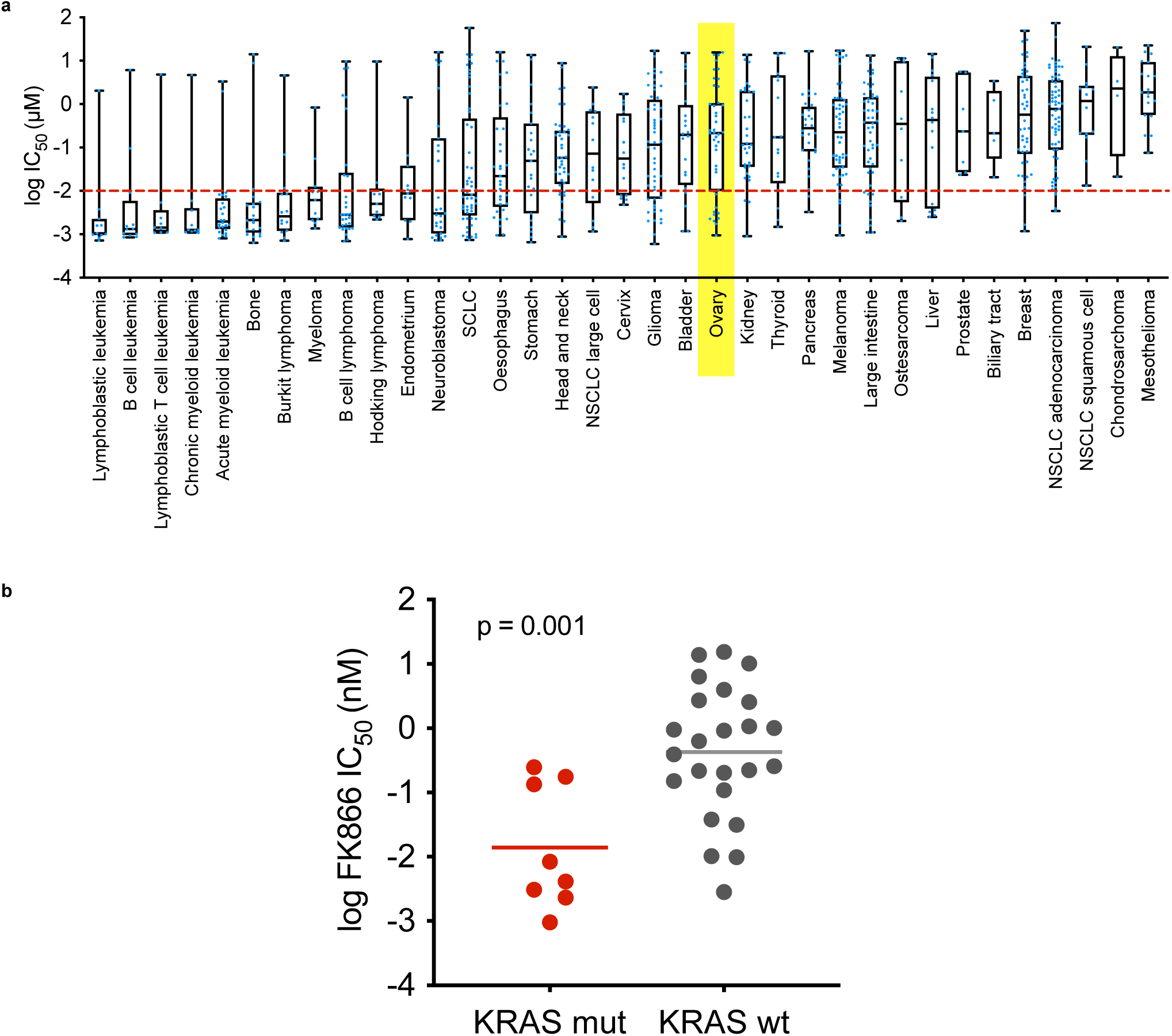
A subset of ovarian tumours are sensitive to the NAMPTi FK866. **a)** Using the Genomics of Drug Sensitivity in Cancer (GDSC) resource (https://www.cancerrxgene.org/) FK866 (NAMPTi) sensitivity was characterised in different cancer cell lines. log IC_50_ values are shown for each cell line in a box and whisker plot, individual points are shown in blue. Ovarian cancer is highlighted in yellow. **b)** The log IC_50_ values observed in *KRAS*-mutant (mut) and -wildtype (wt) EOC cell lines are shown. Statistical significance was determined using the Mann-Whitney U test.

To validate these findings, a panel of ten ovarian cancer cell lines was assembled, with genomic features that reflect different EOC subtypes^26^ (**Fig. 2a**). The panel included six RAS/PI3K-wildtype cell lines. COV318, COV504 and 59M cells have *TP53* mutations. KURAMOCHI and OVSAHO cells possess *TP53* and *BRCA2* mutations, and it should be noted that KURAMOCHI cells are reported to have amplified *KRAS*^26^. COV644 cells do not possess mutations in these pathways. Four RAS/PI3K-mutant cell lines were included in the panel. A2780 cells possess *BRAF*, *PI3K* and *PTEN* mutations. HEY-A8 cells have *BRAF* and *KRAS* mutations. OVCAR-8 cells possess *TP53*, *ERBB2* and *KRAS* mutations, but have been reported to have a heterozygous *BRCA1* methylation^27^. Finally, TOV21G cells have *KRAS*, *PI3K* and *PTEN* mutations. To assess their relative FK866 sensitivity, the panel was treated with the NAMPTi for 6-days and IC_50_ values were calculated (**Fig. 2b and Supplementary Fig. 1a**). This confirmed that the four RAS/PI3K-mutant lines were more sensitive to NAMPT-inhibition than the RAS/PI3K wildtype cell lines.

**Fig. 2:**
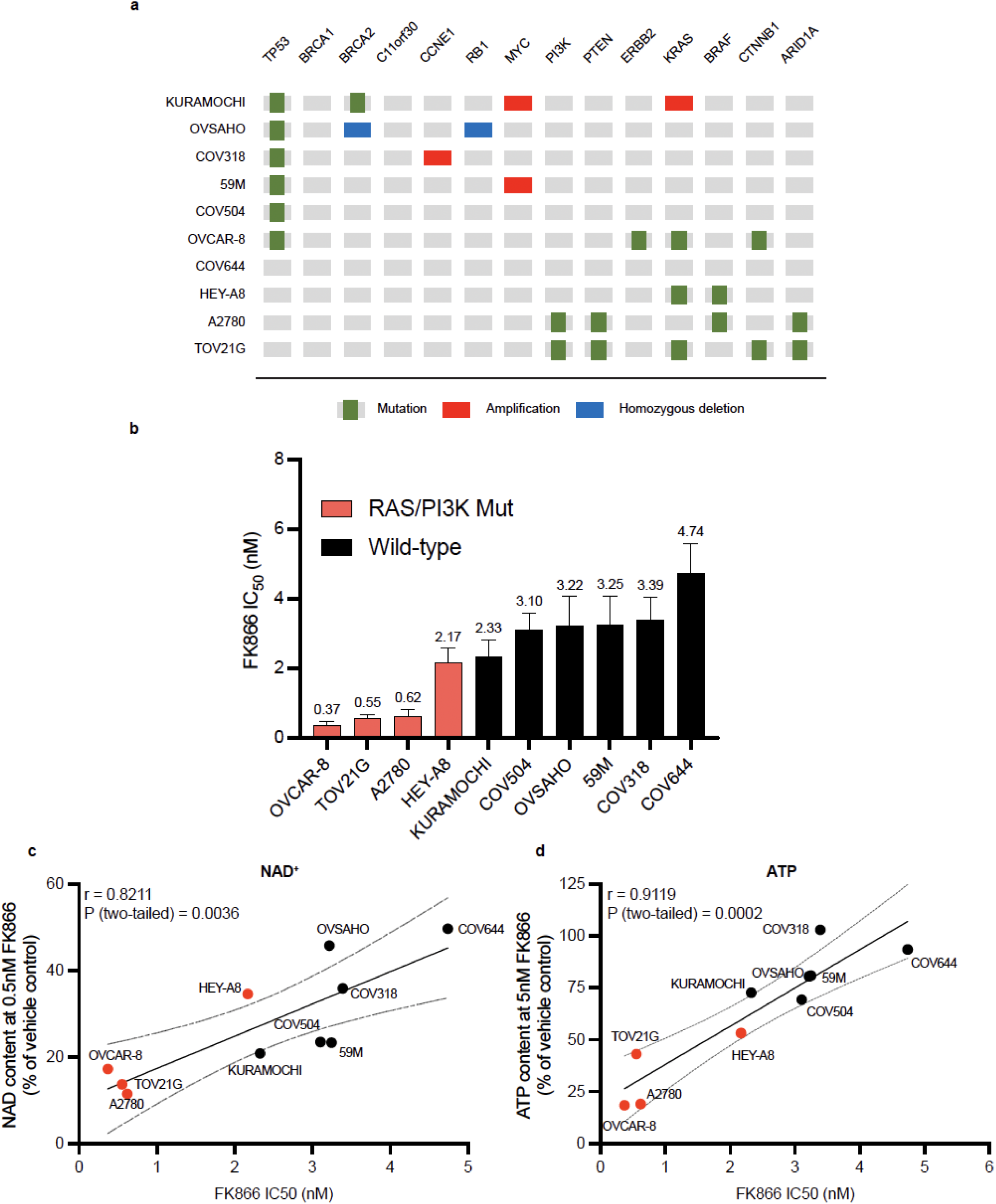
RAS/PI3K pathway mutant EOC cell lines are sensitive to the NAMPT inhibitor FK866. **a)** Mutation profiles of selected epithelial ovarian cancer cell lines, this graphic has been adapted from Domcke et al. (2013)^26^. **b)** EOC cell lines FK866 sensitivity after 6-days treatment. Cell biomass was measured using the SRB assay. IC_50_ values are shown for each cell line, along with their 95% confidence intervals. **c)** NAD^+^ content after 24-hours treatment with 0.5nM of FK866. **d)** ATP content after 48-hours treatment with 5nM of FK866. **c-d)** Data were normalised to cell biomass (SRB assay). Pearson correlation coefficient (R) and linear regression were used to analyse the relationship between NAD^+^/ATP content with FK866 and FK866 sensitivity (IC50). r and p values are shown.

To investigate whether increased FK866 sensitivity in RAS/PI3K-mutant cell lines was due to increased depletion of NAD^+^ pools, cells were treated with FK866 for 24-hours and NAD^+^ assays were performed (**Fig. 2c and Supplementary Fig. 1b**). When treated with a low dose of FK866 (0.5nM) clear differences emerged. The three most sensitive RAS/PI3K mutant cell lines (A2780, OVCAR-8 and TOV21G) achieved the greatest reduction in their NAD^+^ pools (>82% reduction), whilst a ∼65% reduction was achieved in the less-sensitive HEY-A8 cell line. Interestingly, the level of NAD^+^ depletion was found to correlate with FK866 sensitivity (IC_50_) (r = 0.8211, p = 0.0036) (**Fig. 2c**). To gain further insight into the impact of FK866 on NMN and NAD^+^ pools, a UPLC-MS/MS metabolomics assay was performed using COV318 (RAS/PI3K-wildtype) and OVCAR-8 (RAS/PI3K-mutant) cells (**Supplementary Fig. 2a,b**). This revealed that COV318 cells had larger NMN (∼336% larger) and NAD^+^ (∼298% larger) pools when compared to OVCAR-8 cells under basal conditions. Furthermore, 24-hours treatment with 1nM FK866 reduced NMN and NAD^+^ pools to a greater extent in OVCAR-8 cells (91% and 96%, respectively) when compared to COV318 (∼51% and 68%, respectively).

NAMPT inhibition with FK866 inhibits ATP synthesis, which leads to the delayed induction of cell death, as NADH is a substrate for the mitochondrial electron transport chain and NAD^+^ is a co-factor required for glycolysis^28,29^. To investigate the impact of FK866 treatment on ATP levels, the EOC panel was treated for 48-hours with FK866, and an ATP assay was performed (**Fig. 2c and Supplementary Fig. 1c**). This revealed 5nM FK866 could reduce ATP levels in the majority of EOC cell lines tested. However, the greatest reduction were observed in the RAS/PI3K-mutant cell lines (>47%). Interestingly, the level of ATP depletion also correlated with FK866 sensitivity (IC_50_) (r = 0.9119, p = 0.0002) (**Fig. 2c**).

### Combined PARP and NAMPT inhibition is more effective in RAS/PI3K pathway mutant 3D spheroids

We evaluated whether FK866 could enhance PARPi responses in the EOC panel in 2D culture (**Fig. 3a and Supplementary Fig. 3**). In this experiment, cells were treated for 6-days with olaparib and low doses of FK866 (100pM or 500pM) that did not impact cell proliferation as monotherapy. Due to their increased FK866-sensitivity A2780, OVCAR-8 and TOV21G cells were only co-treated with 100pM FK866. In all cell lines tested co-treatment with FK866 reduced olaparib IC_50_ values. Since 3D models better recapitulate the complexity of the *in vivo* tumour microenvironment^30^, COV318 (**Fig. 3b and Supplementary Fig. 4a**), A2780 (**Fig. 3c and Supplementary Fig. 4b**), HEY-A8 (**Fig. 3d and Supplementary Fig. 4c**) and OVCAR-8 cells (**Fig. 3e and Supplementary Fig. 4d**) growing in 3D were treated for 6-days with olaparib and/or FK866, before assessing cell viability. TOV21G cells were not included as they did not form spheroids. Interestingly, co-treatment with FK866 significantly improved PARPi responses in all three RAS/PI3K-mutant cell lines, but not in the RAS/PI3K-wildtype COV318 cell line.

**Fig. 3:**
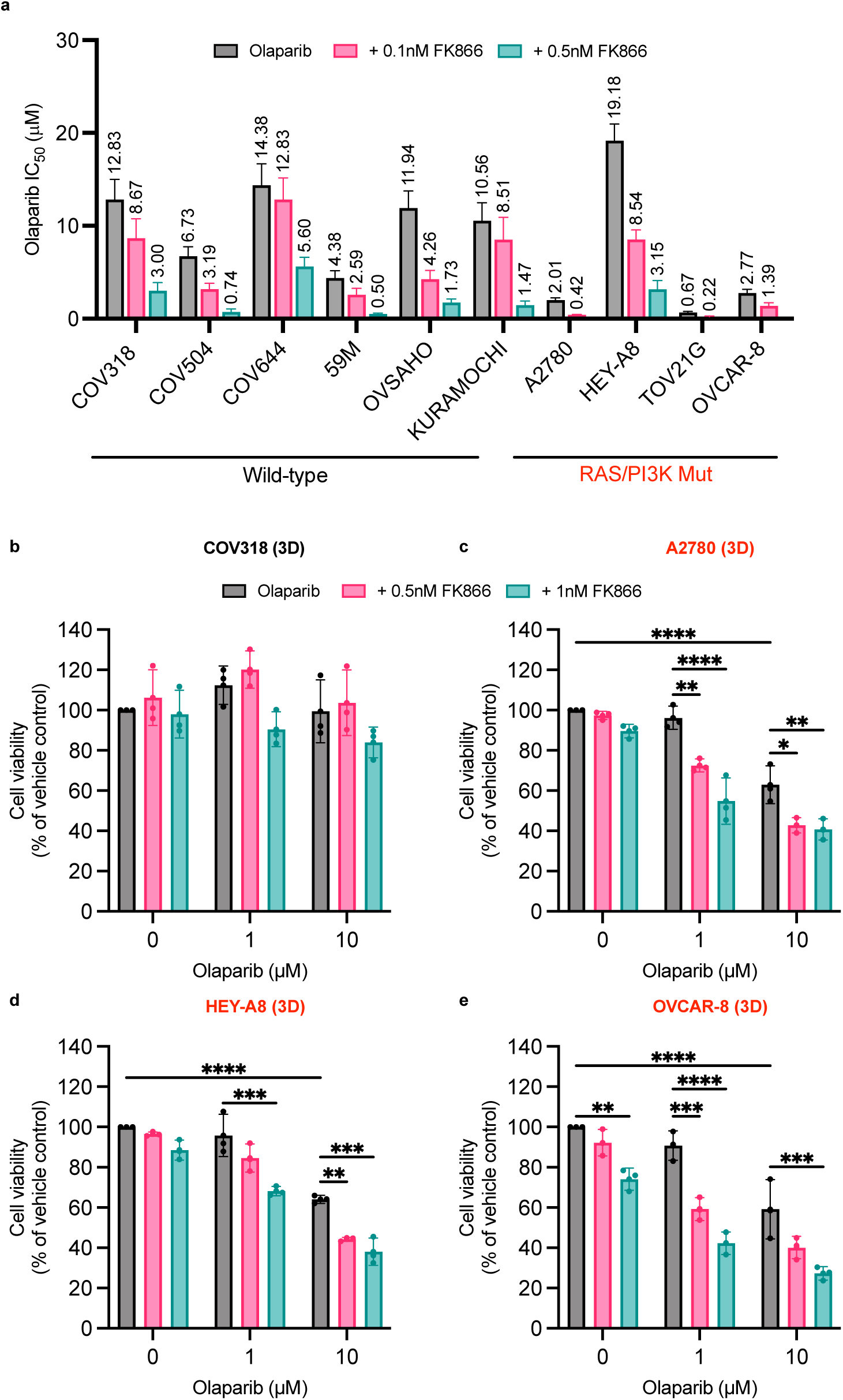
FK866 potentiates the cytotoxic effects of olaparib in EOC cell lines but is more effective in RAS/PI3K pathway mutant 3D spheroids. **a)** EOC cell lines were co-treated in 2D with olaparib and FK866 (0.1nM or 0.5nM) to assess their sensitivity to the combination after 6-days treatment. Cell biomass was measured using the SRB assay. IC_50_ values are shown for each cell line, along with their 95% confidence intervals. **b-e)** 3D spheroids were co-treated with olaparib (1μM or 10μM) and FK866 (0.5nM or 1nM) to assess their sensitivity to the combination after 6-days treatment. Cell viability was measured using the CellTitre-Glo® 3D assay. Data are the average ± SD of three independent experiments. Statistical significance was determined using 2-way ANOVA followed by Turkey’s multiple comparisons test (p < 0.05, **p < 0.01, *** p < 0.001, **** p < 0.0001).

Since inhibiting PARP activity is known to decrease NAD^+^ consumption^31^, we also evaluated the influence of the PARPi/NAMPTi combination on NMN and NAD^+^ pools using a UPLC-MS/MS metabolomics assay (**Supplementary Fig. 2a,b**). This revealed that 24-hours treatment with 1μM olaparib leads to a slight increase in NMN and NAD^+^ levels in COV318 but not in OVCAR-8 cells. However, when cells were co-treated with olaparib and FK866, the NAMPTi-induced reduction of NMN/NAD^+^ pools far outweighed any reduction in NMN/NAD^+^ consumption caused by the inhibition of PARP activity (ADP-ribosylation).

### Combined PARP and NAMPT inhibition induces apoptosis by depleting NMN and NAD^+^ in RAS/PI3K pathway mutant EOC cell lines

Next, we assessed the influence of olaparib, FK866 and the combination on apoptosis induction in COV318 (**Fig. 4a and Supplementary Fig. 5b**), A2780 (**Fig. 4b and Supplementary Fig. 5c**) and TOV21G cells (**Fig. 4c and Supplementary Fig. 5d**). Whether assessing apoptosis/necrosis using Annexin V staining (24-hours) or a caspase 3/7 activation assay (48-hours), FK866 and/or olaparib had little-to-no influence on apoptosis in the RAS/PI3K-wildtype COV318 cell line. However, in RAS/PI3K-mutant A2780 and TOV21G cells, a significant increase in apoptosis/necrosis (**Supplementary Fig. 5c,d**) and caspase activity (**Fig. 4b,c**) could be observed following treatment with the combination. To confirm that FK866 potentiates PARP-inhibitor responses by depleting the NMN/NAD^+^ pool, a rescue experiment was performed, where medium was supplemented with NMN during drug treatment and caspase 3/7 activity was evaluated (**Fig. 4d,e**). As expected, supplementing medium with NMN rescued A2780 cells treated with the combination (**Fig. 4e**), and caspase 3/7 activity was comparable to cells treated with olaparib alone.

**Fig. 4:**
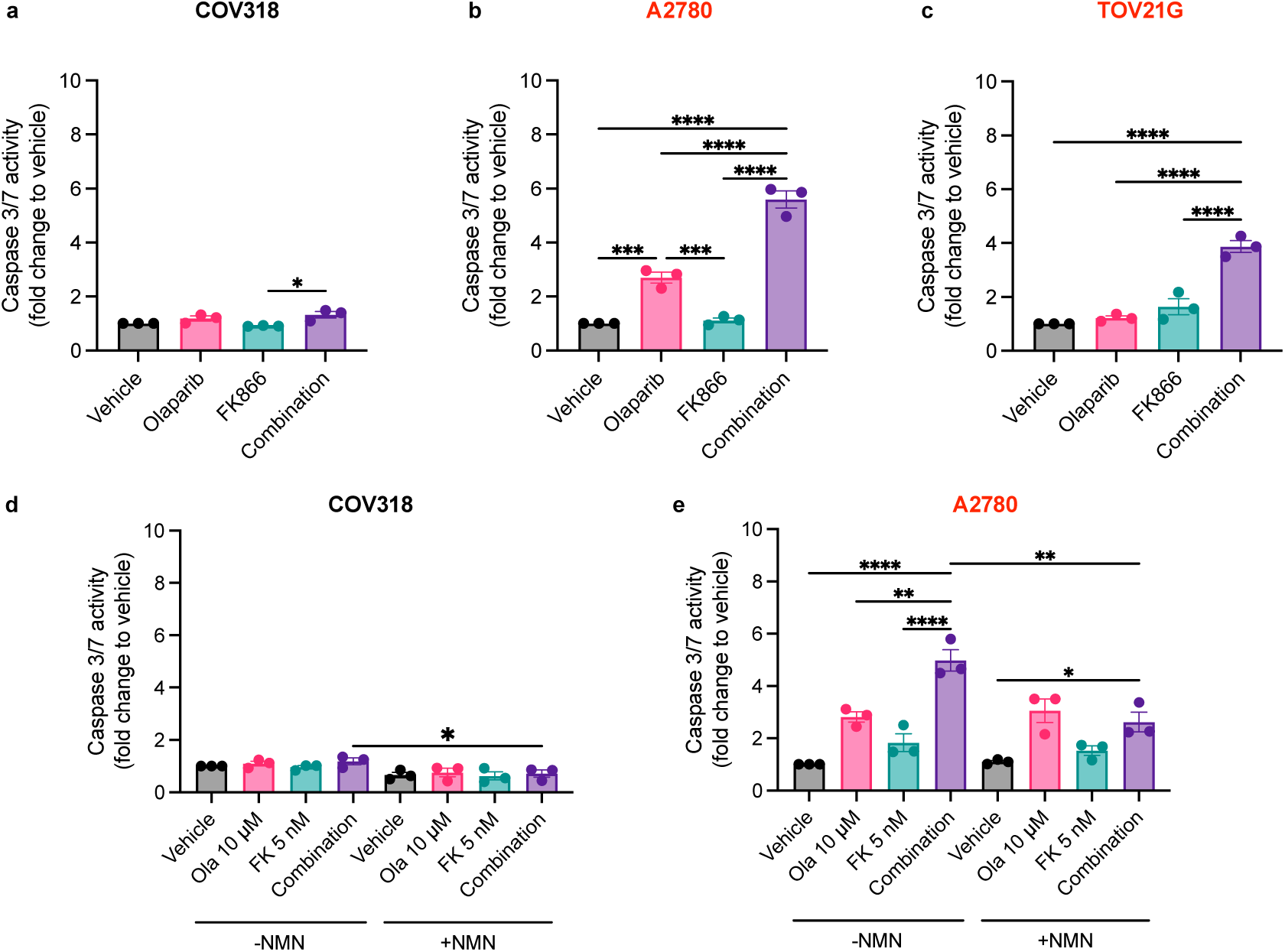
Combination treatment upregulates caspase 3/7 activity in RAS/PI3K pathway mutant EOC cell lines. **a-c)** COV318, A2780 and TOV21G cells were co-treated for 48-hours with olaparib (10μM) and FK866 (5nM) to assess their influence on caspase 3/7 activity (apoptosis). This was measured using the Caspase-Glo® 3/7 assay, data was normalised to cell biomass (SRB assay). **d-e)** A caspase-rescue experiment was performed in COV318 and A2780 cells by supplementing medium with 250μM NMN. Data are the average ± SD of three independent experiments. Statistical significance was determined using 2-way ANOVA followed by Turkey’s multiple comparisons test (p < 0.05, **p < 0.01, *** p < 0.001, **** p < 0.0001).

### ROS-generation promotes the induction of apoptosis following combined PARP and NAMPT inhibition in RAS/PI3K pathway mutant EOC cell lines

Since increased reactive oxygen species (ROS) generation and loss of mitochondrial membrane potential (MMP) is often observed in FK866-induced cell death^28,29^, we evaluated the influence of combined PARP and NAMPT inhibition on ROS levels and MMP. Olaparib and FK866 monotherapy had very little influence on ROS levels in COV318 (RAS/PI3K-wildtype) (**Fig. 5a**) and A2780 (RAS/PI3K-mutant) (**Fig. 5b**) cells. However, when A2780 cells were treated with the combination, a significant increase in ROS could be observed. Both monotherapy and combination treatment had no influence on MMP in COV318 cells (**Fig. 5c**). However, in A2780 cells, loss of MMP was observed in both conditions (**Fig. 5d**). The largest loss of MMP was observed in A2780 cells treated with the combination (4.5-fold increase), and this was followed by cells treated with FK866 (3.8-fold increase) and olaparib alone (2.7-fold increase). To evaluate the influence of ROS generation on apoptosis/necrosis-induction (**Fig. 5e and Supplementary Fig. 5a**) and caspase 3/7 activity (**Fig. 5f**), rescue experiments were performed in A2780 cells, where medium was supplemented with the ROS-scavenger *N*-acetylcysteine (NAC)^32^. Interestingly, NAC supplementation rescues apoptosis/necrosis (**Fig. 5e**), but the increased caspase 3/7 activity observed following combination-treatment was only partially rescued (**Fig. 5f**).

**Fig. 5:**
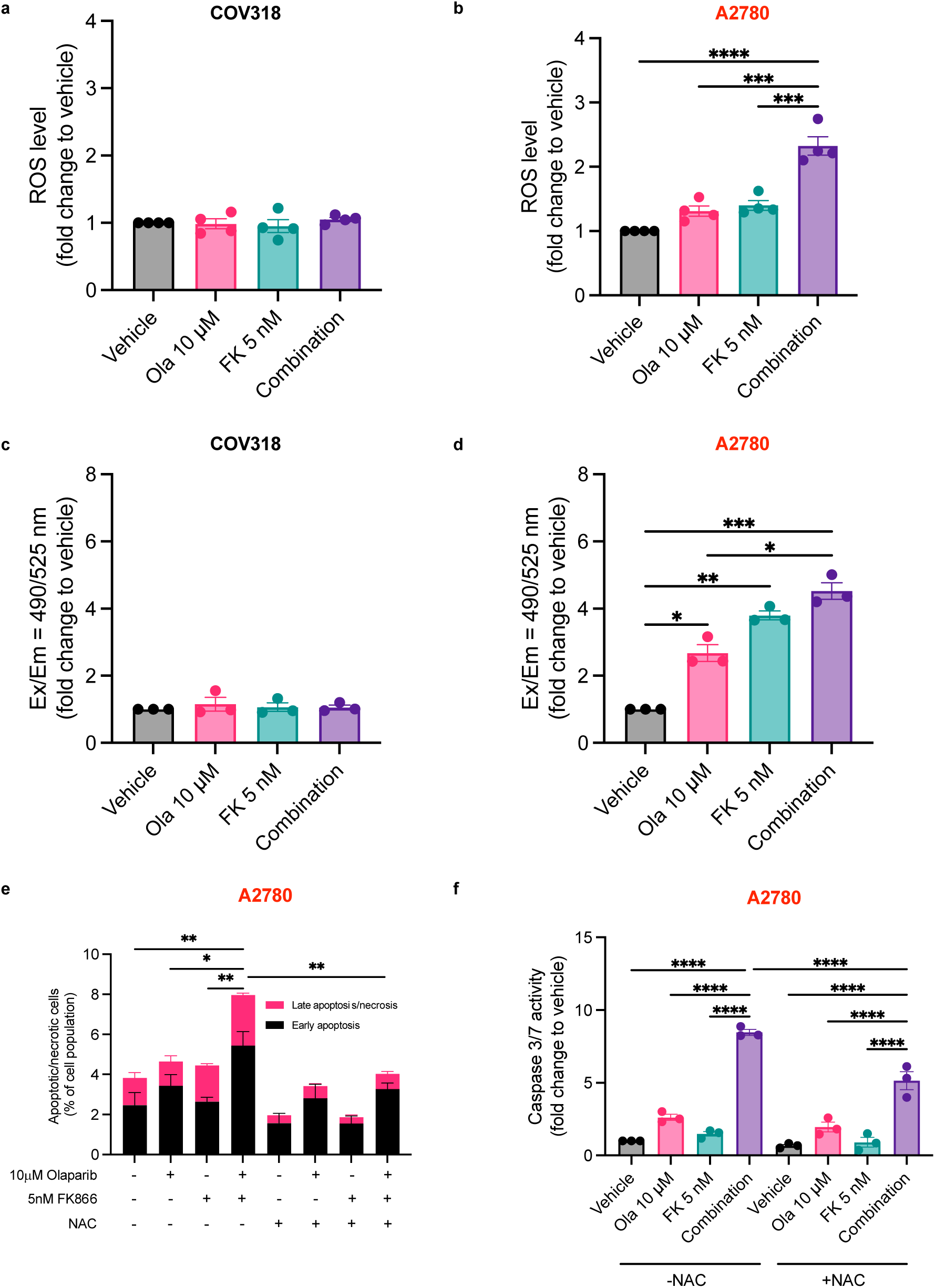
Combination treatment increases ROS generation and mitochondrial dysfunction in RAS/PI3K pathway mutant A2780 cells. **a-b)** COV318 and A2780 cells were treated for 24-hours with the combination and ROS generation was assessed using the ROS-Glo™ H_2_O_2_ Assay. **c-d)** Mitochondrial membrane potential was also assessed after 24-hours treatment using the JC-10 assay. Increases indicate mitochondrial membrane depolarisation. **e)** A2780 cells were treated for 24-hours with vehicle, olaparib, FK866 or the combination before measuring the level of apoptosis/necrosis. As shown in **Supplementary Fig. 5a** live (Annexin V^−^ and propidium iodide^−^ (PI)), early apoptosis (Annexin V^+^ and PI^−^) and late apoptosis/necrosis (Annexin V^+^ and PI^+^) events were quantified. Data was generated using the Annexin V apoptosis assay. **f)** A caspase-rescue experiment was performed in A2780 cells following 48-hours treatment with the combination. Data were generated using the Caspase-Glo® 3/7 assay. All data were normalised to cell biomass (SRB assay). **e-f)** Where indicated, medium was supplemented with 5-10mM of the ROS scavenger NAC (see Methods section for further details). Data represent the average ± SD of at least three independent experiments. Statistical significance was determined using 2-way ANOVA followed by Turkey’s multiple comparisons test (p < 0.05, **p < 0.01, *** p < 0.001, **** p < 0.0001).

### Combined PARP and NAMPT inhibition upregulates DNA damage in RAS/PI3K pathway mutant EOC cell lines

Since the combination has previously been shown to induce apoptosis by upregulating DNA damage^9–11^, we next evaluated the influence of the olaparib/FK866 combination on γH2AX foci formation (**Fig. 6a**). In COV318 cells (RAS/PI3K-wildtype) olaparib and/or FK866 treatment had very little influence on γH2AX foci (**Fig. 6b**). Whereas, in TOV21G cells (RAS/PI3K-mutant) combination treatment led to a significant increase in the number of γH2AX foci (**Fig. 6c**).

**Fig. 6:**
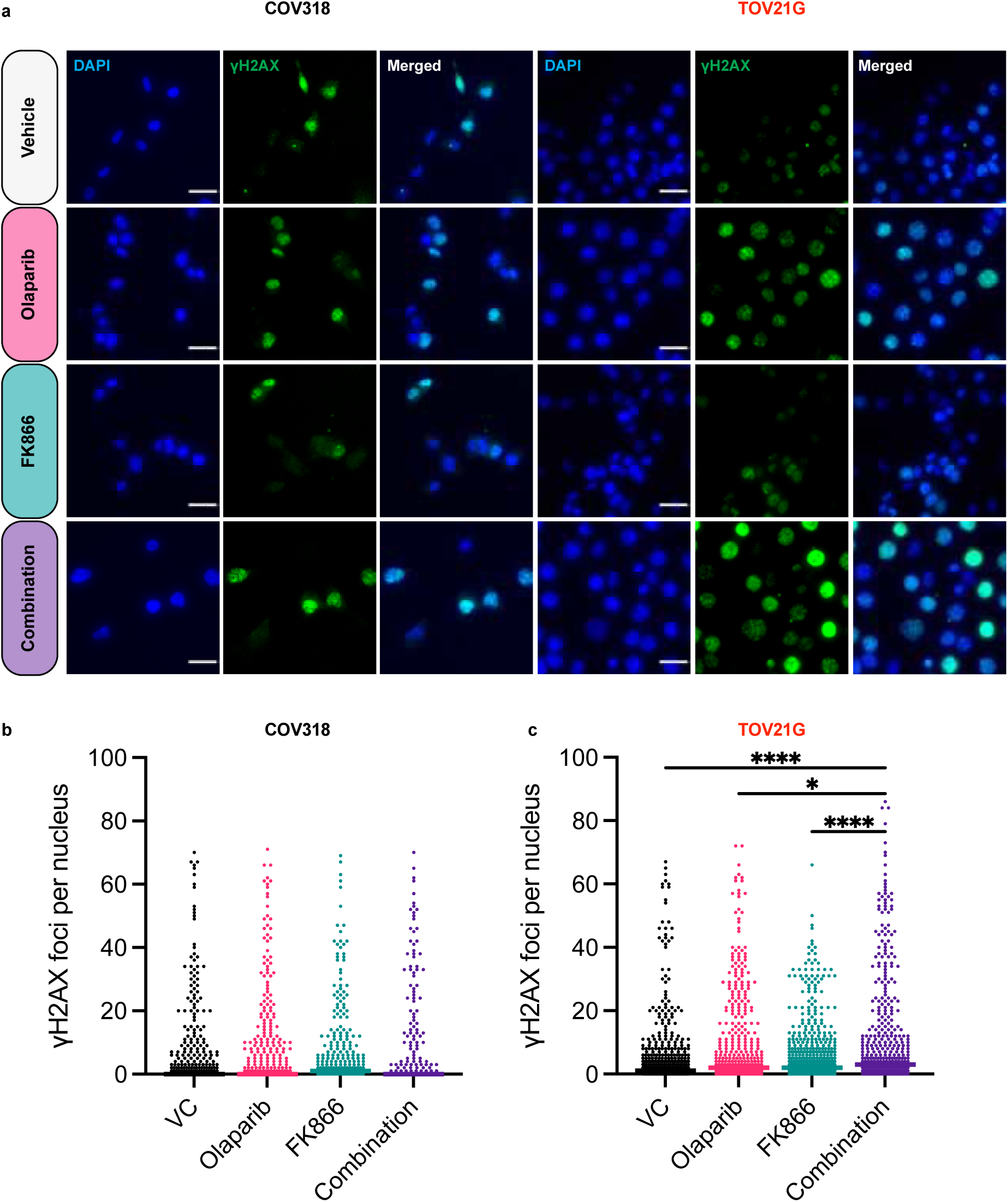
Combined olaparib and FK866 treatment increases γH2AX foci formation in RAS/PI3K pathway mutant TOV21G cells. **a)** Immunofluorescence images of COV318 and TOV21G cells treated for 24hr with 10μM olaparib and/or 5nM of FK866, a 30μm scale bar is shown. **b-c)** Quantitation of γH2AX foci in COV318 and TOV21G cells. Values derived from 400 cells and the median is shown. Data is representative of at least two independent experiments. Statistical significance was determined using 2-way ANOVA followed by Turkey’s multiple comparisons test (p < 0.05, **p < 0.01, *** p < 0.001, **** p < 0.0001).

Given NAMPTi^23,33^ and PARPi^34^ can induce G2/M arrest as single agents, we set out to evaluate the influence of the combination on the cell cycle (**Supplementary Fig. 6**). In COV318 cells (**Supplementary Fig. 6b**), olaparib and/or FK866 treatment had no influence on the proportion of cells in the G2/M phase. Whereas, in A2780 (**Supplementary Fig. 6c**) and TOV21G (**Supplementary 6d**) cells, both monotherapy and combination treatment increased the proportion of cells in the G2/M phase, the latter induced the largest increase.

### Combined PARP and NAMPT inhibition is effective *in vitro* and *in vivo* in the ID8-*Trp53^-/-^_;_ Pten^-/-^* model

Since HGSC is the most common subtype of EOC^17^ and alterations in RAS/PI3K-signalling^14^, such as *PTEN* loss/mutations^35^, may worsen PARPi-responses in the clinic, we investigated whether the combination of a PARP and NAMPT inhibitor would have activity in a transplantable murine model of HGSC with a *Pten* deletion^25^ (i.e., the ID8-*Trp53^-/-^; Pten^-/-^* model). The combination induced a significant decrease in cell biomass when compared to Importantly, combination treatment led to significant upregulation of caspase 3/7 activity (**Fig. 7b**), ROS levels (**Fig. 7c**) and γH2AX foci formation (**Fig. 7d**).

**Fig. 7:**
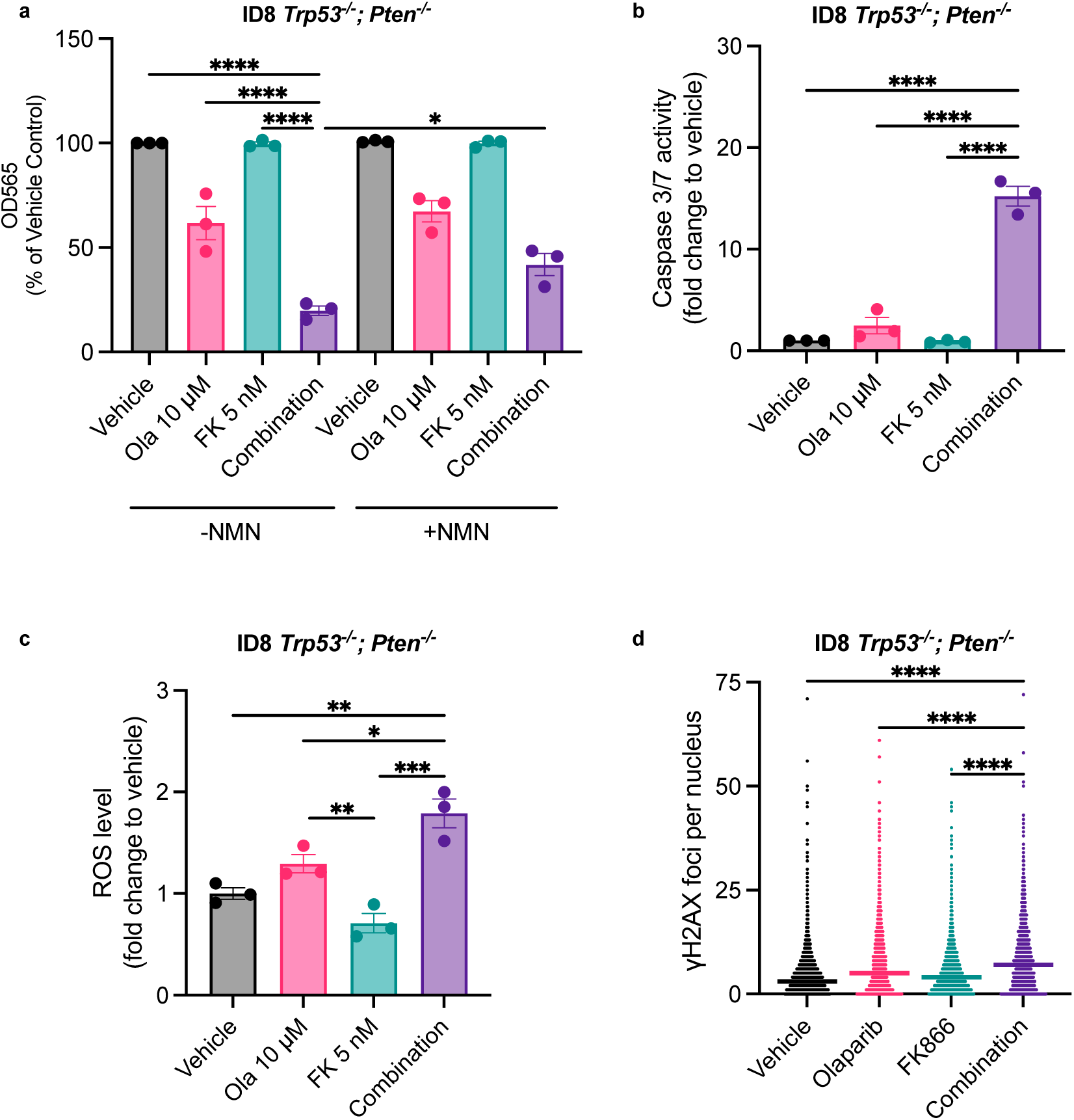
The combination has activity in ID8 *Trp53*^-/-^; *Pten*^-/-^ cells *in vitro*. **a)** ID8 cells were co-treated with olaparib and FK866 (± 250μM NMN) for 3-days. Cell biomass was measured using the SRB assay. The influence of the combination (10μM olaparib and 5nM FK866) on **b)** caspase 3/7 activity (48-hours) **c)** ROS-generation (48-hours) and **d)** γH2AX foci formation (24-hours) was also assessed. Values are derived from 400 cells and the median is shown, data are representative of three independent experiments. **a-c)** Data represent the average ± SD of three independent experiments. Statistical significance was determined using 2-way ANOVA followed by Turkey’s multiple comparisons test (p < 0.05, **p < 0.01, *** p < 0.001, **** p < 0.0001).

We then evaluated the combination *in vivo*. C57BL/6J mice were injected with ID8*-Trp53^-/-^; Pten^-/-^* cells and treated with 10-doses (2-day drug holiday between doses 5-6) of vehicle, FK866 (10mg/kg), olaparib (50mg/kg) or the combination. Treatment commenced after the formation of omental tumours (∼14 days) (**Fig. 8a**). 24 hours after the final injection, there was a significant decrease in omental tumour weight observed in mice that received the combination (**Fig. 8b**). NAD^+^ levels were also significantly reduced following treatment with FK866 or the combination (**Fig. 8c**). Importantly, no significant reduction in mouse weight was observed in any condition (**Supplementary Fig. 7**) indicating that the combination is tolerable.

**Fig. 8:**
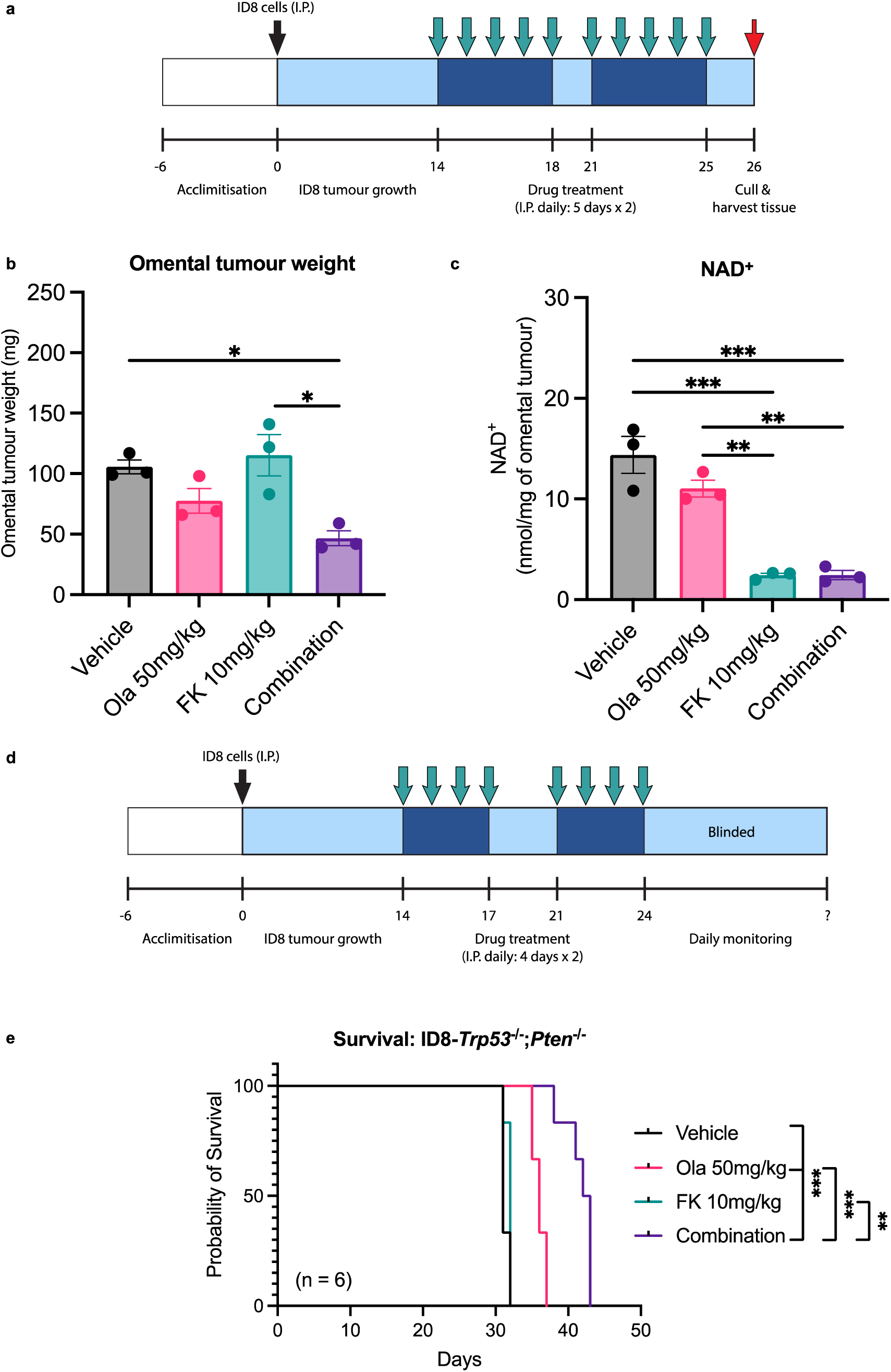
Combined olaparib and FK866 treatment reduces omental tumour weight and improves survival outcomes in the ID8-*Trp53*^-/-^; *Pten*^-/-^ model. **a)** For the endpoint experiment ID8-*Trp53*^-/-^;*Pten*^-/-^ cells were injected intraperitoneally (I.P.) into C57BL/6J mice and allowed to grow for 14-days before mice were treated with vehicle, olaparib, FK866 or the combination. 24-hours after the final drug treatment, omental tumours were **b)** weighed and **c)** harvested for NAD^+^ quantification by LC-MS/MS. Data represent the average ± SD of three independent tumours. Statistical significance was determined using 2-way ANOVA followed by Turkey’s multiple comparisons test (p < 0.05, **p < 0.01, *** p < 0.001, **** p < 0.0001). **d)** For the survival experiment mice, were randomised into different groups for drug treatment. Researchers were blinded after the final drug treatment until the study’s completion for unbiased determination of humane endpoint. **e)** For Kaplan-Meier curves, the log-rank (Mantel-Cox) test was used to determine statistical differences between survival curves. Bonferroni correction was used for the multiple comparisons test (p < 0.05, **p < 0.01, *** p < 0.001, **** p < 0.0001).

Finally, we assessed the effect on survival (**Fig. 8d**). A similar dosing strategy was used, but mice received 8-doses (3-day drug holiday between doses 4-5) of FK866 (10mg/kg), olaparib (50mg/kg), the combination or vehicle. Mice in the vehicle group had a median survival of 31-days, and this only increased to 32-days in mice treated with FK866 (p = NS). Olaparib treatment did provide a moderate survival benefit compared to the vehicle group, increasing median survival to 36 days (p = 0.0006). However, the combination provided the greatest survival benefit, with a median survival of 42.5 days. This was a significant improvement compared to the vehicle (p = 0.0006) and olaparib (p = 0.0007) groups (**Fig. 8e**). Importantly, there was again no significant reduction in mouse weight in any condition (**Supplementary Fig. 8**).

## DISCUSSION

A growing body of evidence has emerged demonstrating that combining NAMPT and PARP inhibitors could be a viable strategy to improve the efficacy of PARPis in diseases such as ovarian cancer^9–11^. However, because dose limiting toxicity has been observed in clinical trials of NAMPTis^12,13^, we set out to identify novel determinants of sensitivity to NAMPTis and the combination.

In this study, we provide the first evidence that EOC cell lines harbouring RAS/PI3K pathway mutations are sensitive to NAMPT-inhibition with FK866. In these cell lines, lower doses of FK866 are required for a critical reduction in NAD^+^ and ATP production to be observed (**Fig. 2b-d**), leading to NAMPTi-induced cell death^28,29^. Previous studies have shown *KRAS*-mutant tumours have lower levels of intracellular NAD^+19,20^, presumably due to increased metabolic demand. Indeed, using LC-MS/MS-based metabolomics we confirmed that the *KRAS*-mutant OVCAR-8 cell line possesses significantly smaller NMN/NAD^+^ pools, that are more susceptible to NAMPT-inhibition when compared to the RAS/PI3K-wildtype COV318 cell line (**Supplementary Fig. 2a,b**). Interestingly, *BRAF^V600E^*mutant melanoma models are known to become dependent on the salvage pathway to synthesise NAD^+^, and are sensitive to NAMPTis^22,23,36,37^, suggesting oncogenic activation of RAS/PI3K-signalling may lead to a NAMPT-dependency in more tumour types. There is evidence that suggests NAMPT-dependency in *RAS*-mutant tumours could be due to regulation of its expression by STAT5 (via the BRAF/ERK pathway)^36,37^. However, the PI3K/AKT-axis may also regulate NAMPT-dependency, as its expression has been shown to inversely correlate with PTEN^21^. This highlights oncogenic activation of Ras-Raf-MEK-ERK and PI3K-AKT-mTOR pathways can lead to the development of an actionable metabolic bottleneck in the NAD^+^ salvage pathway. However, future studies should characterise how different mutations in these pathways can lead to a NAMPT-dependency.

The combination of FK866 and olaparib has previously been shown to be synergistic in ovarian cancer, including in models that are intrinsically resistant to PARPis^11^, as was also observed in our 2D combination experiments (**Fig. 3a**). However, when assessing the combinations influence on cell viability in 3D (**Fig. 3b-d**) or on apoptosis (**Fig. 4 a-c and Supplementary Fig. 5**) we demonstrate for the first time that RAS/PI3K-mutant EOC cell lines benefit to a greater extent following treatment with the combination. Importantly, we confirmed that synergy is dependent on NMN/NAD^+^ depletion (**Fig. 4d,e**), which is in line with previous observations^9–11^.

Previous studies have demonstrated that the on-target activity of FK866 activates apoptotic signalling by inducing mitochondrial dysfunction. Maintenance of MMP is essential to maintain the election transport chain to generate ATP. NAD^+^ plays an important role in this process, its depletion (and NADH consequently) will limit the mitochondrial respiratory substrates availability, leading to a loss of MMP^28,29^. Indeed, our data demonstrate that in the RAS/PI3K-mutant A2780 cell line, FK866-treatment leads to a loss of MMP (**Fig. 5c,d**). To our surprise, given PARP inhibition is reported to lead to mitochondrial protection^38^, we did observe a significant loss of MMP compared to the vehicle group following olaparib treatment (**Fig. 5d**). Furthermore, combined olaparib and FK866 treatment led to the largest MMP loss, although this was not statistically significant when compared to FK866-monotherapy. Generation of ROS is another indicator of defective electron transport in mitochondria, and FK866-induced NAD^+^ depletion has previously been shown to increase ROS levels^28,29^. However, we only observed a significant increase in ROS levels in the RAS/PI3K-mutant A2780 cells following treatment with the combination (**Fig. 5d**). Importantly, using the ROS-scavenger NAC^32^, we demonstrated that it is possible to partially rescue apoptosis following combination treatment in A2780 cells (**Fig. 5e,f**), confirming increased ROS generation upregulates apoptosis. Taken together, these data provide the first evidence that ROS/MMP levels can be altered following combination treatment and contribute towards the induction of apoptosis, further extending our understanding of the mechanism of action of this combination.

The combination has previously been shown to upregulate DNA damage (i.e., increased γH2AX foci)^9–11^, and encouragingly we provide the first evidence that γH2AX foci formation is increased in RAS/PI3K-mutant TOV21G cells (**Fig. 6**). Treatment with a PARPi, such as olaparib, is also associated with marked G2/M cell cycle arrest^34^. Furthermore, treatment with FK866 has also been shown to cause a G2/M-phase arrest-like state^23^. However, to the best of our knowledge the influence of the combination on the cell cycle has not been investigated. Interestingly, in RAS/PI3K-mutant A2780 and TOV21G cells, we observed a significant increase in the proportion of cells in the G2/M-phase following combination treatment, suggesting it enhances G2/M-phase arrest (**Supplementary Fig. 6**). Given a substantial amount of mitochondrial energy (e.g., ATP) is required for cell-cycle progression, especially at the G2/M phase^23,39^, it is possible the combination blocks cells in the G2/M phase since mitochondrial energy production is severely impaired (e.g., ATP loss, MMP loss and increased ROS) in RAS/PI3K pathway mutant cells.

Our combination experiments in ID8 *Trp53^-/-^; Pten^-/-^* cells confirmed that similar trends were observed, namely, growth inhibition that could be rescued by NMN (**Fig. 7a**), and a significant increase in apoptosis (**Fig. 7b**), ROS (**Fig. 7c**) and γH2AX foci (**Fig. 7d**) when compared to monotherapy groups. Finally, our *in vivo* data demonstrate for the first time the efficacy of combined olaparib and FK866 treatment in inhibiting the growth of a *Pten*-null tumour (**Fig. 8**).

In future, it would be interesting to investigate whether targeting other parts of the NAD^+^ biosynthesis pathway could sensitise RAS/PI3K-mutants to PARPis. For example, the Preiss-Handler pathway could be targeted by inhibiting nicotinate phosphoribosyltransferase (NAPRT)^40^. Furthermore, the therapeutic window could be further maximised through the use of antibody-drug conjugates (ADC) to improve the delivery of NAMPTis to cancer cells, whilst sparing healthy tissue^41^. Encouragingly, NAMPTi-ADCs have shown potent anti-cancer activity *in _vivo42-44_*.

One limitation of our study is that ID8 cells derive the ovarian surface epithelium rather than the fallopian tube, which is the origin of most HGSC cases^45,46^. Furthermore, it would be advantageous to validate our findings in genetically engineered murine models that better represent other ovarian cancer subtypes^47,48^, and also in patient-derived models. It should also be noted, the A2780 cell line has previously been misclassified as HGSC, but recent reports demonstrate it is a model of endometroid ovarian cancer, as originally described^26,49^.

Nonetheless, taken together, our data suggest that RAS/PI3K pathway mutations sensitise EOC cells to NAMPT inhibition, increase the therapeutic window of the PARPi/NAMPTi-combination, and could extend the benefit of PARPis to a broader range of EOC patients (e.g., those without a *BRCA1/2* mutation). Given RAS/PI3K pathway mutations are observed in each EOC subtype and are associated with poorer PARPi-responses, we suggest that combining PARPis and NAMPTis may be a promising strategy in this group of EOC patients. Especially with the development of new generations of NAMPTi and PARPi, such as OT-82^33,50–52^ (currently in clinical trials: NCT03921879) and Saruparib^53,54^ (clinical trial: NCT04644068), with more favourable toxicological profiles, further expanding the possibilities for PARPi/NAMPTi combination therapy.

## ACKNOWLEDGEMENTS

M.G., E.W.T., and H.C.K. acknowledge funding from the EPSRC (EP/S023518/1) and Cancer Research UK Convergence Science Centre (CANTAC721\100021); Y.M. from an Imperial College London and China Scholarship Council joint PhD studentship (202208310101); Yi.X. from an Imperial College London President’s PhD Scholarship; A.N. from an AstraZeneca/NIHR Imperial BRC Imperial College Research Fellowship; A.B. and A.S. from NIHR Imperial BRC; F.M. and J.B. from a STRATiGRAD and MRC DTP PhD studentship (MR/N014103/1); C.Bray. from a MRC iCASE Enterprise DTP PhD studentship (MR/R015732/1); Ovarian Cancer Action (A.B., S.S., K.T., A.N., C.W. and I.M.). I.M. also acknowledges support from an NIHR Senior Investigator Award. We thank animal technicians from the Imperial College London CBS animal centre for supporting animal studies.

## AUTHOR CONTRIBUTIONS

Conceptualisation: M.G., A.B., I.M., and H.C.K.; methodology: M.G., S.S., Yi.X., Cr.B., A.B., and H.C.K.; Investigation: M.G., Yi.X., L.A., Y.M. Cr.B., K.T., S.S., Y.X., F.M., Ch.B., and A.B.; visualisation: M.G., Yi.X., L.A., Y.M., and A.B.; funding acquisition: A.B., E.W.T., and H.C.K.; project administration: A.B., E.W.T., and H.C.K.; supervision: A.B., E.W.T., and H.C.K.; writing original draft: M.G. All authors contributed to the review and editing of the paper.

## DECLARATION OF INTERESTS

E.W.T. is a past Director, founder and shareholder in Myricx Bio and consults for and/or receives research funding from Kura Oncology, Pfizer, Samsara Therapeutics, Myricx Pharma, Merck Sharp and Dohme (MSD), Exscientia, Dunad Therapeutics and Daiichi Sankyo. A.B. is currently a paid employee of and has ownership interests at GlaxoSmithKline; however, contributions to this work were made during their employment at Imperial College London.

## METHODS

### Cell culture

All human cell lines were cultured in phenol red-free DMEM (ThermoFisher, A1443001) supplemented with 5.6mM glucose (ThermoFisher, A2494001), 10% foetal bovine serum (FBS; ThermoFisher, 10270106), 2mM L-glutamine (ThermoFisher, 25030-081), 100 units/ mL penicillin, 100μg/mL streptomycin (ThermoFisher, 15070-063). Cells were maintained at 37°C in a humidified atmosphere containing 10% CO_2_. Cell lines were authenticated using Short Tandem Repeat (STR) profiling by Public Health England. To avoid genetic drift human cell lines were cultured for no more than ten passages. Cell lines were regularly tested for mycoplasma contamination.

The generation of ID8*-Trp53^-/-^; Pten^-/-^* cells has been described previously^25^. For *in vitro* experiments, these cells were adapted to physiological glucose (5.6mM) over the course of three passages. After adaptation, cells were cultured in phenol red-free DMEM supplemented with 5.6mM glucose, 10% foetal bovine serum, 2mM L-glutamine, 100 units/mL penicillin, 100μg/mL streptomycin and 1mM sodium pyruvate. For *in vivo* experiments, cells were cultured as described previously^24,25^, in phenol red-free DMEM supplemented with 25mM glucose, 4% FBS, 2mM L-glutamine, 100 units/mL penicillin, 100μg/mL streptomycin, 1mM sodium pyruvate (ThermoFisher, 11360070) and 1x Insulin-Transferrin-Selenium (ThermoFisher, 41400045).

### Annexin V and PI assay

COV318, A2780 and TOV21G cells were seeded into 6-well plates and allowed to attach overnight. The next day, medium was replaced with DMEM supplemented with 0.1% DMSO, FK866 and/or olaparib. After 24-hours, cell surface annexin V expression was quantified as described previously^55^.

### ATP assay

Cell lines were seeded onto two parallel plates at a density of 10,000 cells per well. 24-hours after seeding cells were treated with FK866. After 48-hours treatment, SRB plates were processed using the SRB assay protocol, and white walled plates were processed using the CellTiter-Glo® assay, following manufacturers guidelines. Luminescence was measured on a FLUOstar Omega Microplate reader (BMG LABTECH). Cell free wells were used as blanks to remove background luminescence. ATP were normalised to cell biomass (SRB assay).

### Bioinformatics

Cell sensitivity data (IC_50_) to FK866 (Daporinad) was downloaded from the Genomics of Drug Sensitivity in Cancer web portal (www.cancerrxgene.org, GDSC1). HEY-A8, OVCAR-8, OVCAR-5, OVK-18, TOV21G, OV-7, SW26, OV-56 cell lines were labelled as KRAS mutant^26^.

### Caspase-Glo^®^ 3/7 assay

24 hours after seeding, cells were treated with vehicle, olaparib and/or FK866. After 2-days treatment, SRB plates were processed using the SRB assay protocol, and white walled plates were processed using the Caspase-Glo^®^ 3/7 Assay, following manufacturers guidelines. Luminescence was measured on a FLUOstar CLARIOstar Plus Microplate Reader (BMG LABTECH). Cell free wells were used as blanks to remove background luminescence. Caspase data was normalised to cell biomass. Data was then normalised to VC wells.

### Cell cycle assay

COV318, A2780 and TOV21G cells were seeded into 6-well plates and allowed to attach overnight. The next day, medium was replaced with DMEM supplemented with 0.1% DMSO, FK866 and/or olaparib. Cell cycle assay was performed as described previously^55^. Briefly, after 24-hours drug treatment, cells were incubated with 200 mM IdU (Sigma, I7125) and washed with PBS twice before collection by trypsinization (Gibco, 12604013). After fixing overnight at 4°C in 70% ethanol, chromosomes were denatured by adding 2M HCl with 0.5% Triton X-100 (Sigma-Aldrich, T8787) with agitation and incubating for 30-minutes. Next, 0.1M of NaB4O7 (pH 8.5; Sigma, B9876) was added to samples, and they were incubated for 5-minutes to neutralise the HCl. The samples were then incubated with 1% BSA (Sigma, A2153) and 0.2% Tween 20 (Sigma, P1379) in PBS for 30-minutes for blocking. To probe IdU, the samples were incubated with mouse anti-IdU antibody (Abcam, ab6326; 1:25 in blocking buffer (BD Bioscience, 347580)) for 60-minutes each at room temperature. The samples were then washed with blocking buffer (BD Bioscience, 347580) twice and incubated with AF488-conjugated goat anti-mouse antibody (Invitrogen, A11001; 1:100 in blocking buffer (BD Bioscience, 347580)) before 15-minutes incubation with the FxCycle PI/RNase staining solution (Invitrogen, F10797) in the dark. The samples were then analysed using BD FACS Canto analyser and data were analysed using FlowJo V10.6.2.

### Immunofluorescence

For immunofluorescence assays, COV318, TOV21G and ID8-*Trp53*^-/-^; *Pten*^-/-^ cells were treated for 24-hours with vehicle, FK866 and/or olaparib. At endpoint, cells were washed with 100μL PBS, before being fixed for 15 minutes (RT) with 50μL 4% PFA (ThermoFisher, 11586711) in PBS. Plates were washed with 100μL PBS and permeabilised for 20 minutes with 50μL 0.1% Triton-X100 (Sigma-Aldrich, T8787) in PBS. After two PBS washes, a blocking step was performed (1% BSA and 2% FBS in PBS) for 30 minutes at RT. 1% BSA (in PBS) alone or with the γH2AX (1:500) antibody (Sigma-Aldrich, 05-636) was added and plates were incubated overnight at 4°C. The next day, wells were washed three times with PBS and once with 1X TBS-T. Next, cells were incubated in the dark (RT) for 2 hours with 50μL 1% BSA (in PBS) staining solution containing: Goat anti-Mouse IgG (H+L) Cross-Adsorbed Secondary Antibody, Alexa Fluor™ 488 (1:500, Invitrogen, #A11001) and 1 μg/mL DAPI (Sigma-Aldrich, #9542). Wells were then washed three times with PBS and one time with 1X TBS-T, before adding 200μL of PBS to each well. Prior to imaging AbsorbMax™ film (Sigma-Aldrich, Z722537) was placed on top of 96-well plates to minimise background fluorescence. Images were then taken on an IN Cell^TM^ Analyser 1000 (GE Healthcare Life Sciences) and the accompanying INCell analyser 3.5 software. A 20x objective lens was used and 25 fields were captured in each well. CellProfiler V4.0.6 (Broad Institute of MIT and Harvard) was used for image analysis.

### *In vivo* experiments

*In vivo* experiments complied with UK welfare guidelines and were conducted under UK Home Office personal and project licence (PA780D61A) authority in dedicated facilities. All experiments were approved by the Imperial College Animal Welfare and Ethics Review Board (reference IM170119). 5×10^6^ ID8-*Trp53*^-/-^; *Pten*^-/-^ cells were injected into 6–8-week-old female C57BL/6 mice by intraperitoneal (I.P.) injection. Mice were monitored daily by observation of weight and overall health. Vehicle solution consisted of 3.5% DMSO and 6.5% Tween 80 (Sigma-Aldrich, P4780) in 0.9% saline. The vehicle solution was used to prepare suspensions of olaparib (50mg/kg) and FK866 (10mg/kg). Each suspension was vigorously vortexed and sonicated to break down particulates, and drug solutions were inspected using a haemocytometer to ensure no large particles remained. Drugs were stored as aliquots at −20°C and thoroughly vortexed prior to use.

Vehicle, olaparib and/or FK866 were administered as 200 µl IP injections. For the endpoint experiment, mice received daily I.P. injections on days (D) 14-18 and D21-25 inclusive. The day after the final injection (D26) mice were killed. Organs/tumour were collected, weighed and snap frozen.

For the survival experiment, mice received daily I.P. injections on D14-17, and D21-24 inclusive. After completion of the dosing schedule, researchers were blinded until the study’s completion for unbiased determination of humane endpoint. Mice were killed upon reaching a humane endpoint. Organs/tumour was collected, weighed and snap frozen.

### MMP assay

COV318, A2780 and TOV21G were seeded in triplicates into black 96-well plates. The cells were also seeded into parallel clear 96-well plates at the same density for normalization against cellularity. After overnight incubation to allow attachment to the plates, medium was replaced with fresh DMEM with 0.1% DMSO, FK866 and/or Olaparib. After 48-hours drug treatment, mitochondrial membrane potential (MMP) was measured using the JC-10 assay kit (Abcam, ab112134) according to the manufacturer’s instructions. In brief, 50μl of assay buffer A with JC-10 dye was added to each well and incubated at room temperature for 1-hour. Next, 50μl of assay buffer B was added to each well to lyse the cells and fluorescence at excitation/emission=490/525nm (cut off at 515nm) was measured by FLUOstar CLARIOstar Plus Microplate Reader (BMG LABTECH). The parallel clear plates were processed according to the SRB assay. Cell-free wells were used for background subtraction. Fluorescent intensity was normalised against SRB data for correction of cell biomass.

### NAD^+^/NADH-Glo^TM^ assay

Cell lines were seeded onto two parallel plates. A clear plastic plate was used for an SRB assay and white walled 96-well plates were used for the NAD/NADH-Glo™ assay (Promega, G9072). 24-hours after seeding, cells were treated with FK866. After 24-hours treatment, SRB plates were processed using the SRB assay protocol, and white walled plates were processed using the NAD^+^/NADH-Glo™ assay. In brief, medium was aspirated from wells and replaced with 50μL PBS, or 50μL PBS with diluted NAD+ standards (50-2000nM) was added into cell free wells. 50μL 1% dodecyltrimethylammonium bromide (DTAB) with 0.2N NaOH was added to each well and plates were briefly mixed to ensure homogeneity and cell lysis. 50μL lysate was transferred into an empty well and 25μL 0.4N HCl was added. Plates were covered with foil and incubated for 15 minutes on a ThermoMixer® C (Eppendorf) heat block at 60°C. Plates were allowed to equilibrate for 10 minutes at RT before adding 25μL 0.5M Trizma® Base (Sigma-Aldrich, 93352) solution. 50μL of this mixture was then transferred into a white walled 96-well plate along with 50μL of the NAD^+^/NADH-Glo detection reagent, which was made up with reagent at the following ratio: 1000μL Luciferin Detection Reagent, 5μL Reductase enzyme, 5μL Reductase substrate, 5μL NAD Cycling Enzyme and 25μL NAD Cycling Substrate. Plates were mixed for 30 minutes at RT before luminescence was measured on a FLUOstar Omega Microplate reader (BMG LABTECH). Cell free wells were used as blanks to remove background luminescence. Data were normalised to VC wells. NAD^+^ levels were determined by normalising to cell biomass (SRB assay).

### ROS assay

COV318, A2780, TOV21G and ID8 *Trp53*^-/-^; *Pten*^-/-^ cells were seeded into white 96-well plates, and a parallel clear 96-well plate was seeded for normalisation against cell biomass (SRB assay). After overnight incubation and cell attachment to the plates, medium was removed and 80μl of fresh DMEM media with 0.1% DMSO, FK866 and/or Olaparib was added to each well. During ROS rescue experiments, 2-hours prior to drug-treatment cells were pre-treated with 10mM NAC. The NAC concentration was reduced to 5mM during drug treatment. After 24- (human) or 48-hours (ID8) H_2_O_2_ was measured using ROS-Glo H_2_O_2_ assay (Promega, G8820) according to manufacturer’s manual. Briefly, 6-hours before the endpoint is reached, 20μL of combined H_2_O_2_ substrate (provided in the kit) and test compound were added to reach the final concentration of H_2_O_2_ substrate at 25μM and drug concentrations were kept unchanged. After a 6-hour incubation, 100μL ROS-Glo™ detection solution (provided in the kit) was added, and the plates were incubated at room temperature for 20-minutes. Next, luminescence was measured using a FLUOstar CLARIOstar Plus Microplate Reader (BMG LABTECH). To subtract background signals, cell-free wells were processed using the same protocol. To correct for cell biomass, luminescence signal was normalised to SRB readings from the parallel plates (SRB assay).

### Sulforhodamine B (SRB) cell growth assay

Human cell lines were seeded on 96-well plates at a pre-determined density. The following day, wells were aspirated and replaced with medium containing vehicle (0.1% DMSO), FK866 (50pM-50nM) (APExBIO, A4381) or olaparib (1nM-200μM) (Selleckchem, S1060). Both inhibitors were prepared in DMSO (Sigma-Aldrich, D2650). Where indicated, drugs were added in combination and/or medium was supplemented with 250μM of NMN (Sigma-Aldrich, N3501). After 3- (ID8) or 6-days (Human) of treatment, cells were fixed for 1 hour at 4°C with 10% (w/v) trichloroacetic acid (TCA) solution (Sigma-Aldrich, T6399). Wells were washed with water and left to dry at room temperature (RT) overnight. Cells were stained for 30 minutes at RT with 0.4% SRB (Sigma-Aldrich, 230162) solution (w/v) in 1% (v/v) acetic acid (ThermoFisher, 10005920). Wells were washed four times with 1% acetic acid (v/v) to remove excess dye and plates were left to dry overnight at RT. 200μL 10mM Tris Base solution (Sigma-Aldrich, T1699) was added to wells and plates were placed on a shaker to solubilise the protein bound SRB dye. Absorbance was then measured on a SpectraMax Gemini XPS Microplate Reader (Molecular Devices) at 565nm. A minimum of three technical replicates were used for each condition. Cell free wells were used as blanks to remove background absorbance. Data were normalised to vehicle control (VC) wells and dose response curves were generated using nonlinear regression function [inhibitor] vs. response (three parameters) in the GraphPad Prism 10.0.0 software.

### Statistics

Unless otherwise stated, statistical tests were performed using GraphPad Prism (10.0.0). p<0.05 was considered significant.

### Targeted LC-MS/MS analysis for NMN and NAD^+^ quantification

#### Cell lines

COV318 and OVCAR-8 cells were seeded on 6 well plates at a density of 200,000 cells per well. The following day, media was aspirated and replaced with fresh media containing vehicle, FK866 and/or olaparib. Separate plates were included for media blanks, extraction blanks and for cell counts at the time of extraction to normalise data by cell number. 24-hours after treatment commenced, cell counts were performed on a Vi-CELL XR Cell Viability Analyzer (Beckman coulter). The remaining plates were transferred onto a ThermalTray^TM^ (Biocision) that had been pre-cooled with dry-ice and 2mL Eppendorf tubes were pre-cooled in a CoolRack^®^. Wells were aspirated and cells were washed with ice-cold Ringers solution (2 mL, one tablet per 500 mL of MilliQ H_2_O, Sigma-Aldrich, 96724). Wells were aspirated and cells were quenched with 1mL of ice cold LC-MS grade 80:20 methanol:water (v/v) (ThermoFisher, 34966) for 20 minutes. Wells were scraped and the extracts were transferred to the 2mL tubes. The previous step was repeated with 500μL 100% methanol. Next, extracts were centrifuged for 20 minutes at 18,000 x g (4°C) using a Eppendorf™ Centrifuge 5430 R (Eppendorf), to pellet out cellular debris. Supernatants were collected in high recovery LCMS vials, dried under nitrogen flow, and stored at −80°C. Before LC-MS/MS analysis samples were reconstituted in 50μL of LC-MS grade H_2_O (ThermoFisher, 10505904), vortexed for 1 minute and stored in an autosampler at 10°C.

LC-MS/MS assays were performed using an Agilent 1290 Infinity LC system (Agilent Technologies) and a 4000 QTRAP triple quadrupole instrument (AB SCIEX) coupled to an ESI source (Turbo V). An ACQUITY UPLC^®^ HSS (High Strength Silica) T3 Column (Waters) was used for Reversed-Phase (RP) Liquid Chromatography, with a column temperature of 40°C. To increase the columns lifetime a ACQUITY UPLC HSS T3 VanGuard Pre-column (Waters Corp, 186003976) was used. Elution was performed at a flow rate of 600μL/min using the gradient described in **Supplementary Table 1** and 5μL of sample was used per injection. Needle wash steps were performed with Acetonitrile:Isopropanol:H_2_O (40:40:20, v/v/v) (ThermoFisher, 10055454) (ThermoFisher, 10684355). Mobile phase A was comprised of H_2_O and 0.2% formic acid (v/v) (Scientific Laboratory Supplies, 56302). Mobile phase B contained acetonitrile (ACN) and 0.2% formic acid (v/v).

To optimise conditions for targeted metabolomics in Multiple Reaction Monitoring (MRM) mode (QqQ) standards for each compound were directly infused into the MS/MS analyser. From this MRM transitions, declustering potential (DP) and collision energy (CE) values were identified for each target analyte as shown in **Supplementary Table 2**.

The Analyst (v1.6.2, SCIEX) software was used to perform peak integration. Purified stock compounds were used to produce a calibration curve (0.025-10μg/mL) for each run. Using this and cell counts, molar concentrations per million cells were calculated for each metabolite.

#### Omental tumours

Roughly 10-15mg of omental tumour was weighed and placed into clean homogeniser tubes containing 0.8mL ice cold 80% Methanol and 0.1mm Precellys® Glass Beads (ThermoFisher, 12927835). Tissue was homogenised using the Precellys Evolution tissue homogeniser (Bertin) with an activated Cryolys Evolution cooling system (18,000 x g, 4 x 20 sec cycle with a 30 sec pause, two cycles). Tubes were centrifuged at 21,000 x g for 10 minutes (4°C). Supernatants were collected in Eppendorf tubes. An additional 800μL 80% Methanol was added, previous steps were repeated, and extracted supernatants for each sample were pooled. Samples were then dried under nitrogen flow and stored at −80°C. A processing blank was also included during the extraction.

Samples were then subjected to dual-phase extraction. 300μL chloroform (ThermoFisher, 390760025)/methanol (2:1) was added to each sample, and tubes were vortexed. Tubes were vortexed and subsequently centrifuged at 21,000 x g for 10 minutes (4°C). Each sample’s aqueous (upper) and organic (lower) layers were transferred into separate LC-MS vials, before drying under nitrogen flow, and storage at −80°C.

On the day of LC-MS/MS assays, omental tumour (aqueous phase) samples were re-suspended in 100μL H_2_O and vortexed for 30 seconds and stored in an autosampler at 10°C. LC-MS/MS assays were performed using an Agilent 1290 Infinity LC system (Agilent Technologies) and a 4000 QTRAP triple quadrupole instrument (AB SCIEX) coupled to an ESI source (Turbo V).

Acquity UPLC ® BEH Amide 1.7μm 2.1 x 150mm Column (Waters, 186004802) was used for HILIC Liquid Chromatography, with a column temperature of 50°C. To increase the columns lifetime a Acquity UPLC ® BEH Amide 1.7μm VanGuardTM Pre-Column 2.1 x 5mm (Waters, 186004799) was used. Elution was performed at a flow rate of 500μL/min using the gradient described in **Supplementary Table 3** and 5μL of sample was used per injection. Needle wash steps were performed with Acetonitrile:H_2_O (1:1, v/v). For HILIC positive mode, mobile phase A was comprised of ACN and 0.1% formic acid (v/v). Mobile phase B contained H_2_O, 20mM ammonium formate (Supelco, 70221) and 0.1% FA. Peak integration was performed as described previously, but for concentration determination, molar concentrations were calculated per mg of omental tumour tissue. MRM transitions, declustering potential (DP) and collision energy (CE) values were identified for each target analyte as shown in **Supplementary Table 4**.

### 3D spheroid viability assay

COV318, A2780, HEY-A8 and OVCAR-8 cells were used for 3D spheroid assays. Cell suspensions were passed through a 0.45μm filter before seeding 100μL of cells onto ultra-low attachment (ULA) 96-well black plates with a clear round bottom, at a density of 5,000 cells per well. Cells were left for 24-hours to allow spheroids to form, and 100μL of media containing either VC or drug (2X concentration) was added to each well. At endpoint, 150μL of media was carefully removed from each well and 50μL of the CellTiter-Glo 3D cell viability assay (Promega, G9682) reagent was added. Cell viability was measured following manufacturers guidelines. Luminescence was measured on a FLUOstar Omega Microplate reader (BMG LABTECH). Cell free wells were used as blanks to remove background luminescence. Data were normalised to VC wells.

## SUPPLEMENTARY FIGURES

**Supplementary Figure. 1:**
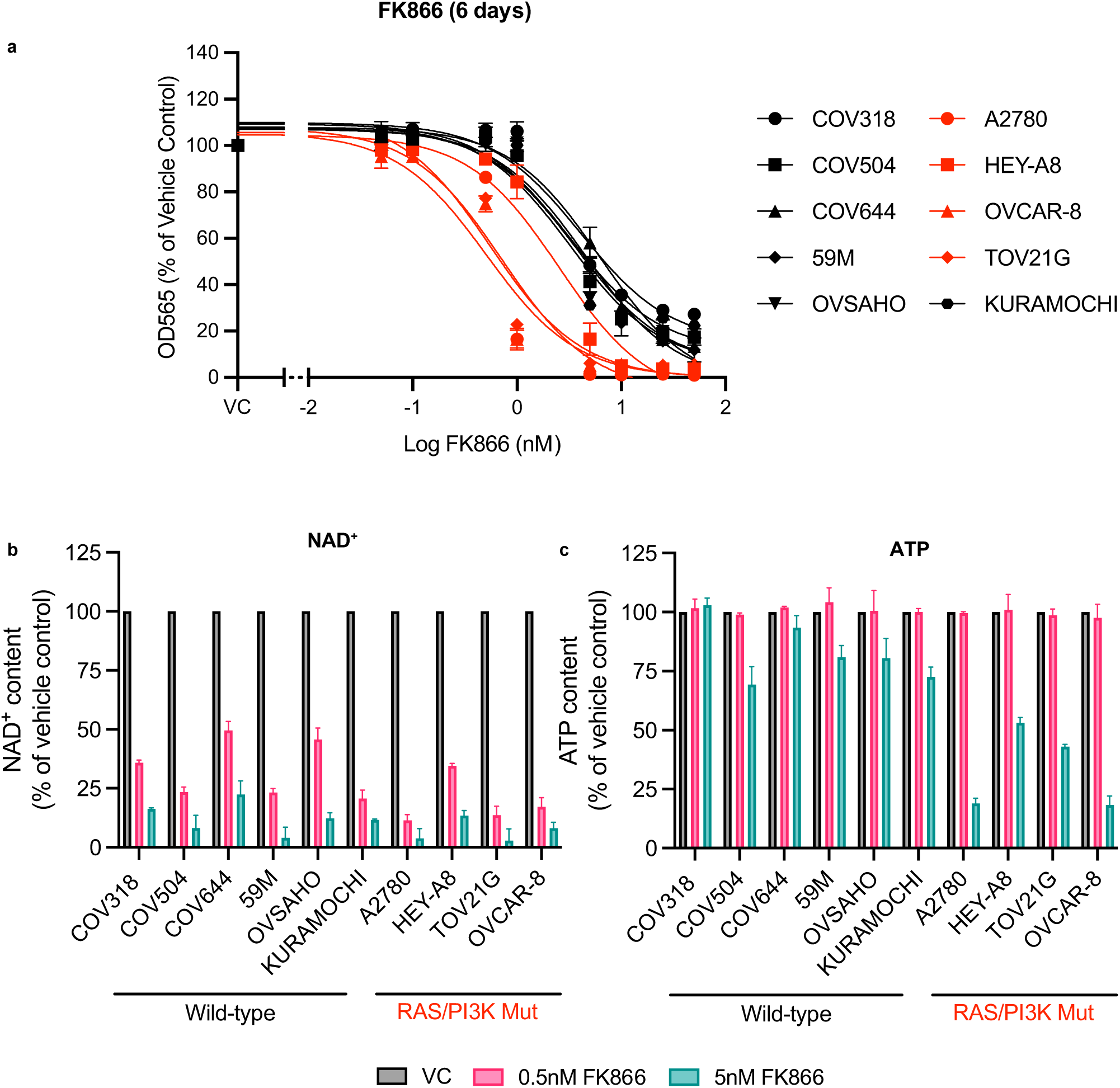
FK866 treatment depletes NAD^+^ and ATP pools to a greater extent in RAS/PI3K-mutant EOC cell lines. **a)** EOC cell lines FK866 sensitivity after 6-days treatment. Cell biomass was measured using the SRB assay. EOC cell lines were treated with vehicle (0.1% DMSO) or FK866 (0.5nM or 5nM) for **b)** 24-hours to measure NAD^+^ content using the NAD/NADH-Glo^TM^ assay or for **c)** 48-hours to measure ATP content using the CellTiter-Glo^®^ 2.0 assay. **b-c)** Data was normalised to cell biomass (SRB assay). Data is the average ± SD of three independent experiments.

**Supplementary Figure. 2:**
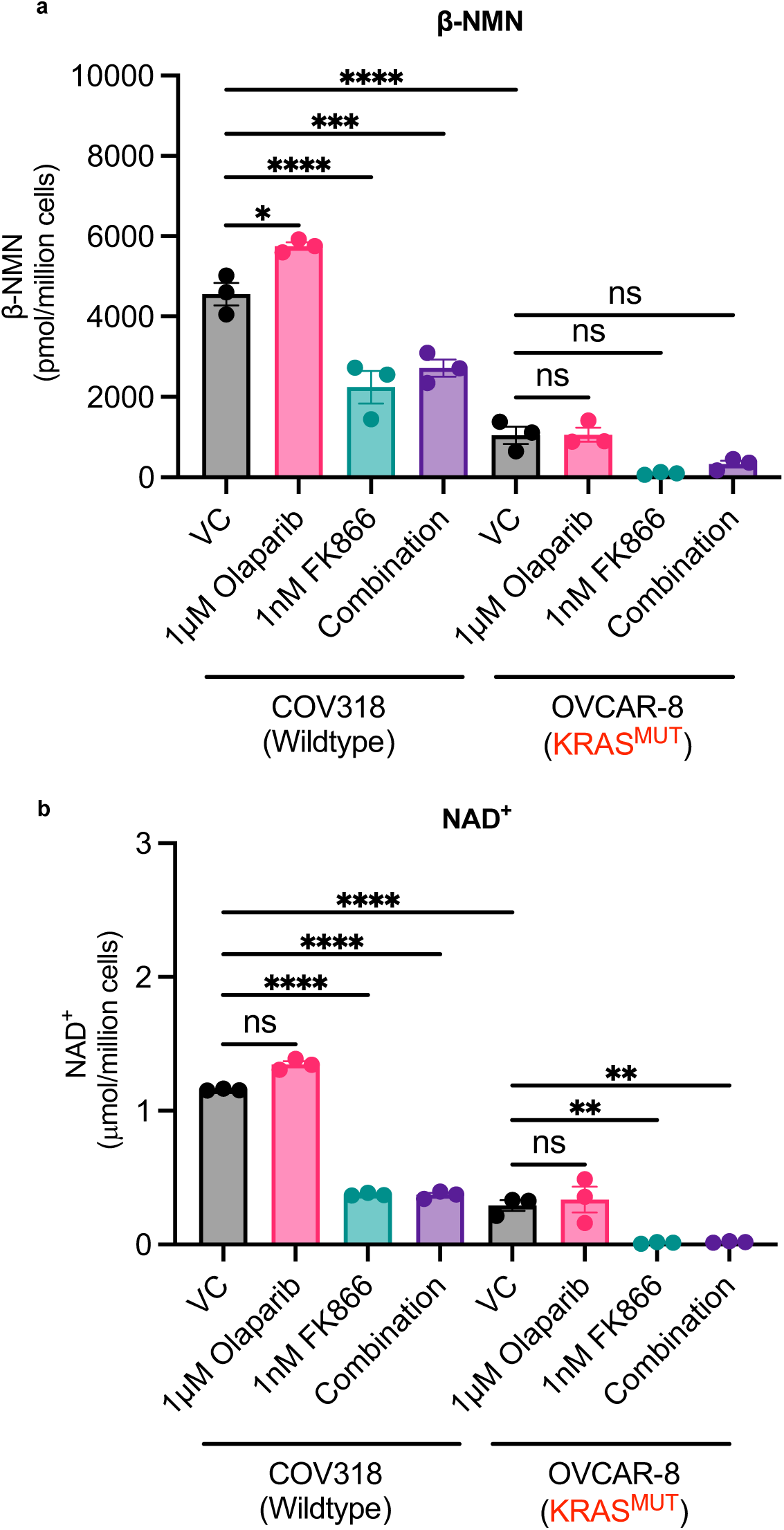
Combined olaparib and FK866 treatment depletes β-NMN and NAD^+^ pools to a greater extent in RAS-mutant OVCAR-8 cells. **a-b)** COV318 and OVCAR-8 cells were treated for 24-hours with vehicle, olaparib, FK866 or the combination, and then samples were extracted for UPLC-MS/MS analysis. The concentration of **a)** β-NMN and **b)** NAD^+^ were calculated using a standard curve. Data is the average ± SD of three independent experiments. Statistical significance was determined using 2-way ANOVA followed by Turkey’s multiple comparisons test (p < 0.05, **p < 0.01, *** p < 0.001, **** p < 0.0001).

**Supplementary Figure. 3:**
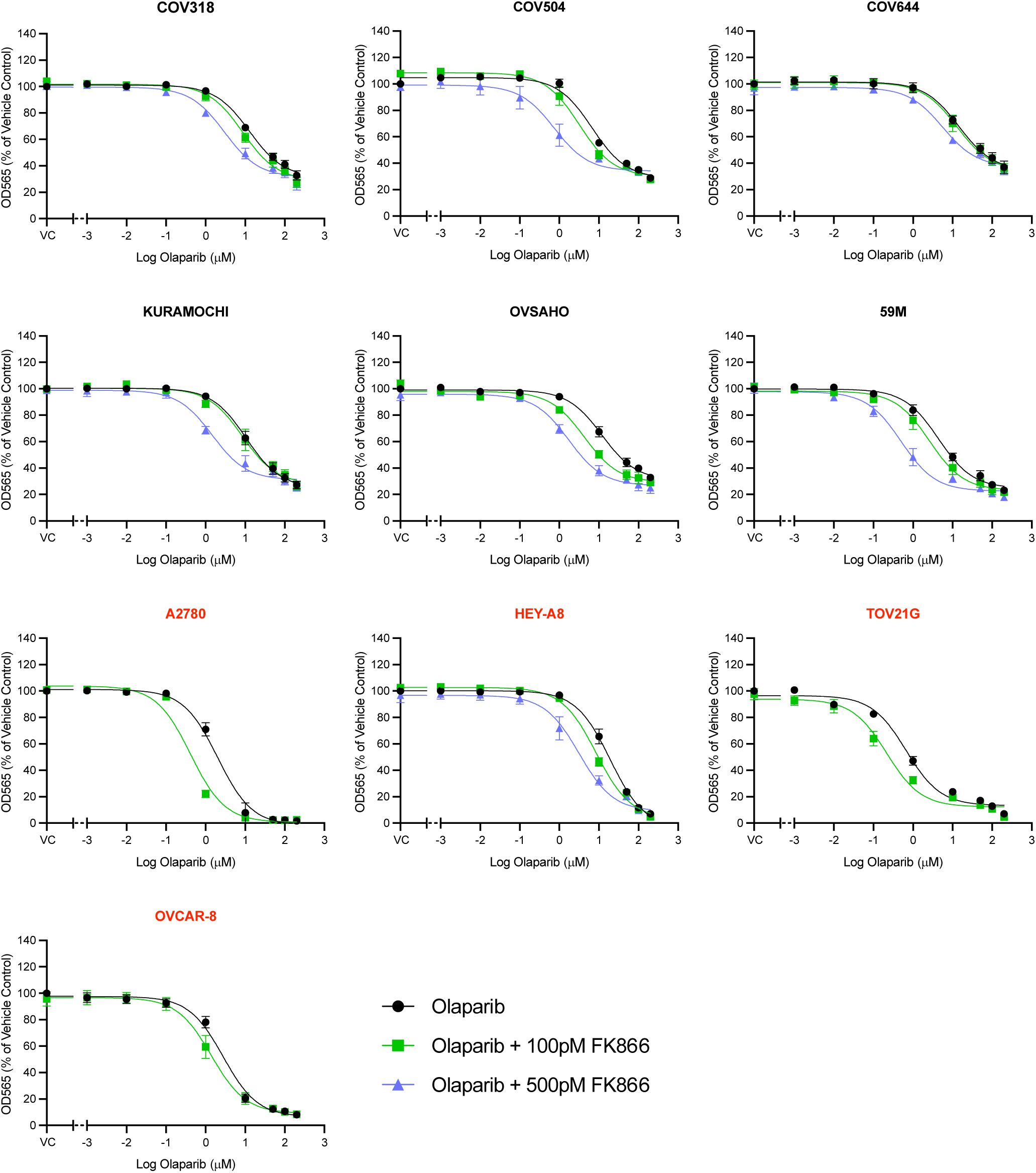
FK866 potentiates the cytotoxic effects of olaparib in EOC cell lines in 2D culture. EOC cell lines were co-treated with olaparib and indicated doses of FK866 for 6-days. Cell biomass was measured using the SRB assay. Data is the average ± SD of three independent experiments.

**Supplementary Figure. 4:**
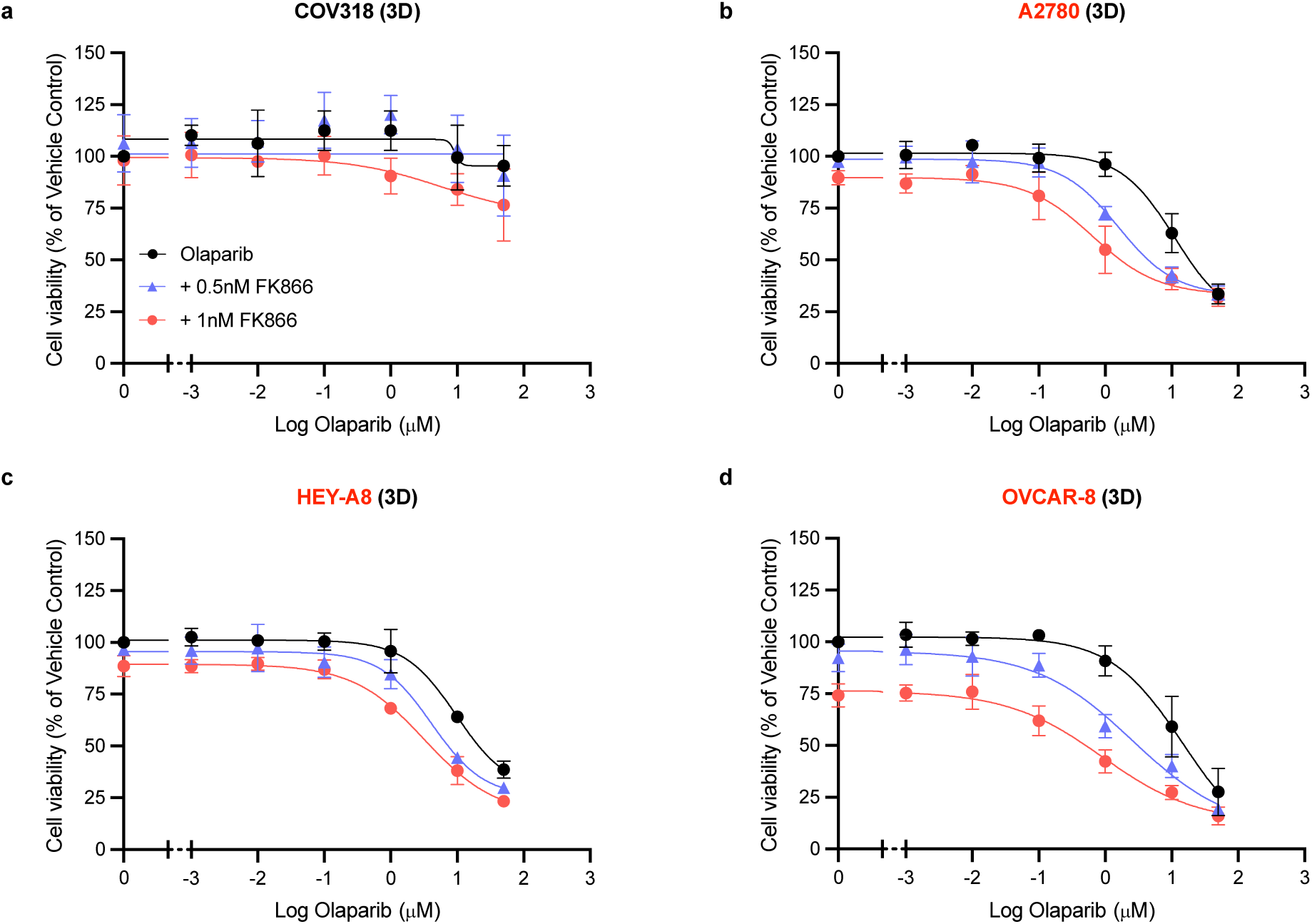
FK866 potentiates the cytotoxic effects of olaparib in RAS/PI3K-mutant EOC cell lines in 3D culture. **a)** COV318, **b)** A2780, **c)** HEY-A8 and **d)** OVCAR-8 spheroids were co-treated with olaparib and FK866 (0.5nM or 1nM) to assess their sensitivity to the combination after 6-days treatment. Cell viability was measured using the CellTitre-Glo® 3D assay. Data are the average ± SD of three independent experiments.

**Supplementary Figure. 5:**
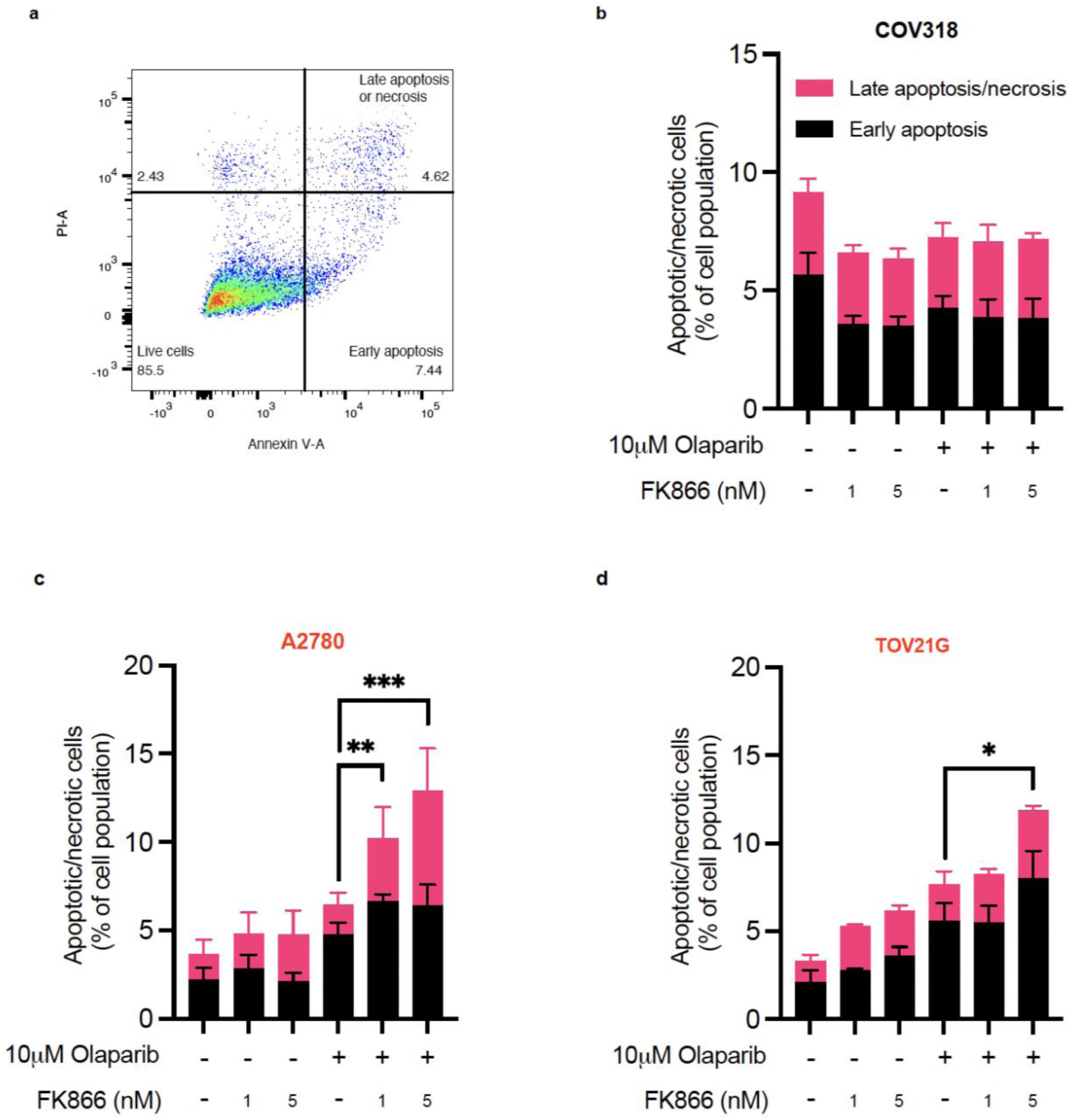
The combination induces apoptosis/necrosis in RAS/PI3K mutant EOC cell lines. **a)** Example of gating used for the Annexin V apoptosis assay. **b)** COV318, **c)** A2780 and **d)** TOV21G cells were treated for 24-hours with vehicle, olaparib, FK866 or the combination before measuring the level of apoptosis/necrosis. Data is the average ± SD of three independent experiments. Statistical significance was determined using 2-way ANOVA followed by Turkey’s multiple comparisons test (p < 0.05, **p < 0.01, *** p < 0.001, **** p < 0.0001).

**Supplementary Figure. 6:**
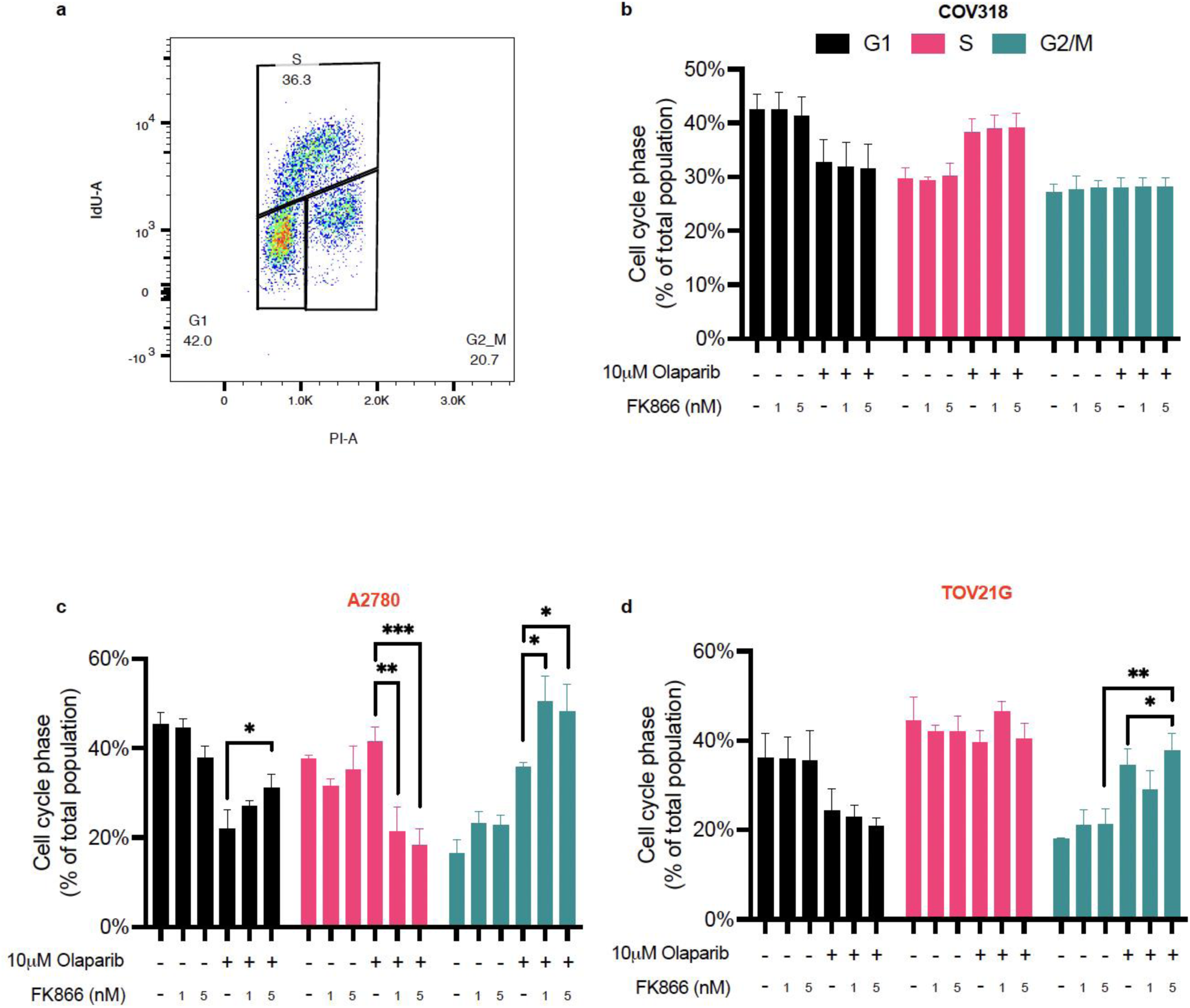
The combination upregulates G2/M arrest in RAS/PI3K-mutant EOC cell lines. **a)** Example gating for the determination of cells in G1, S or G2/M phases by flow cytometry. **b)** COV318, **c)** A2780 and **d)** TOV21G cells were treated for 24-hours with vehicle, olaparib, FK866 or the combination before measuring cell cycle progression. Data is the average ± SD of three independent experiments. Statistical significance was determined using 2-way ANOVA followed by Turkey’s multiple comparisons test (p < 0.05, **p < 0.01, *** p < 0.001, **** p < 0.0001).

**Supplementary Figure. 7:**
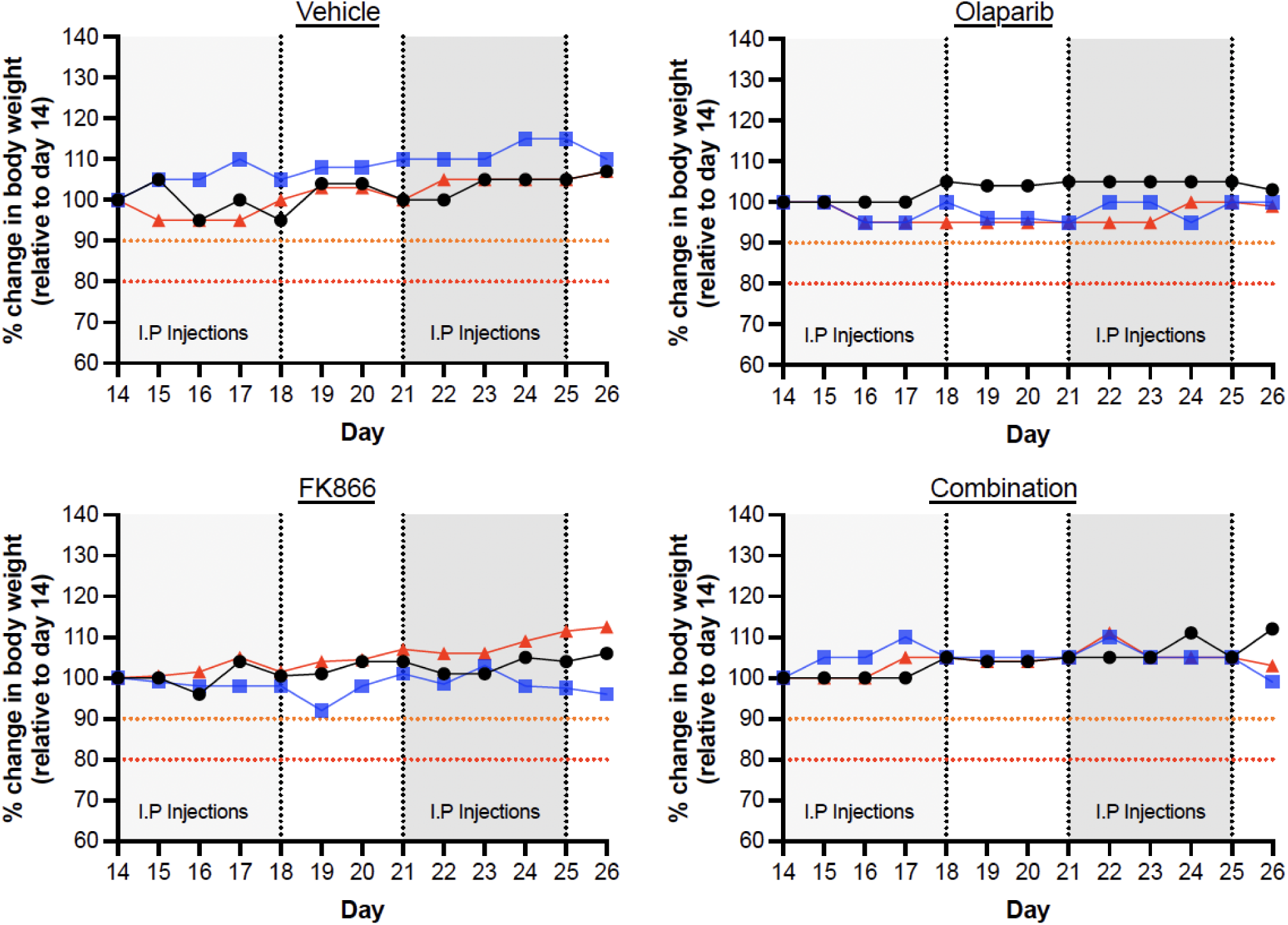
Relative body weight changes of C57BL/6J mice during the *in vivo* endpoint experiment. Relative body weight changes were measured each morning. In each condition data is from three separate mice. Orange and red dotted lines on the y-axis indicate the threshold for 10% and 20% (maximum limit) drops in body weight, respectively.

**Supplementary Figure. 8:**
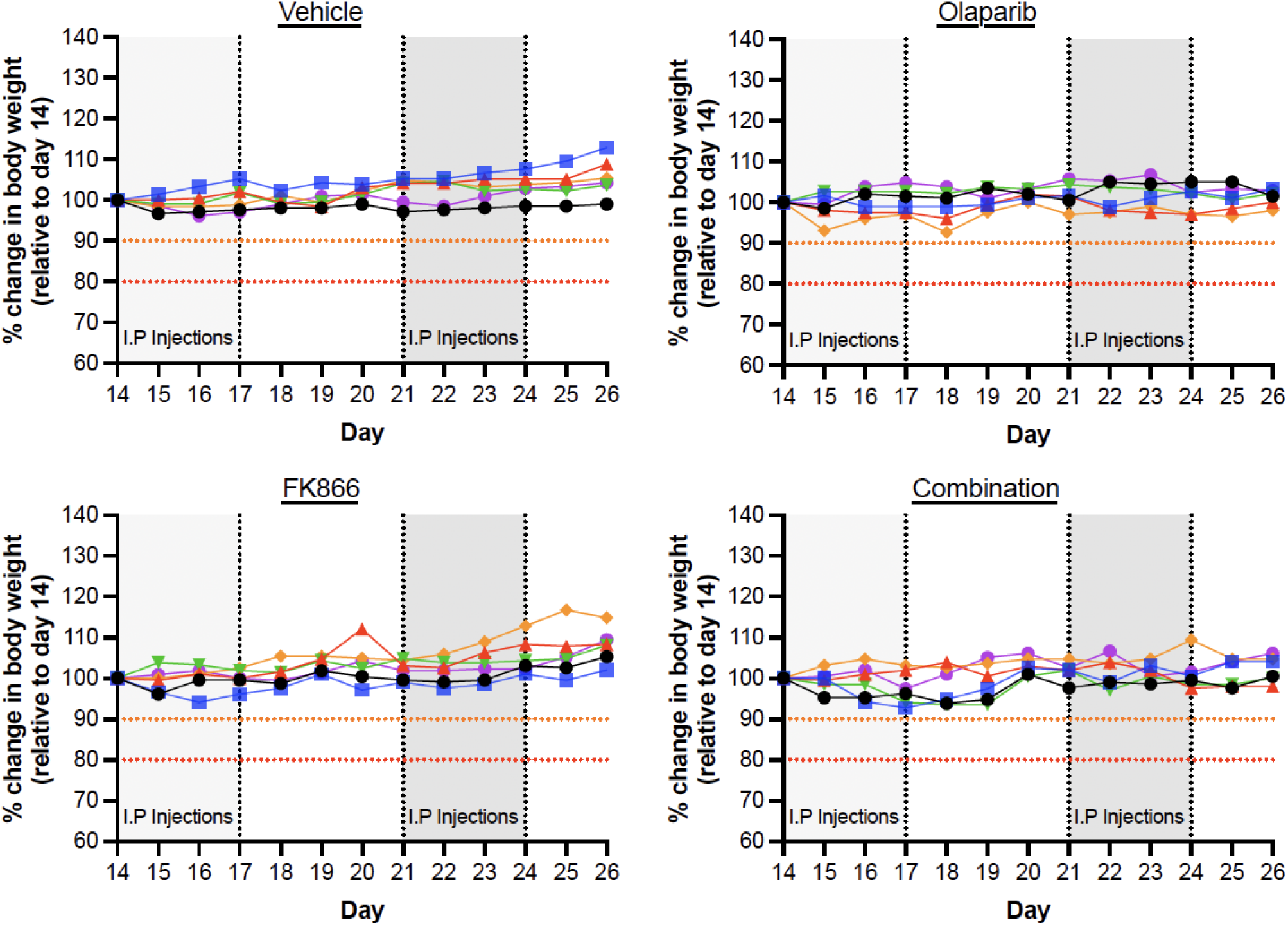
Relative body weight changes of C57BL/6J mice during the *in vivo* survival experiment. Relative body weight changes were measured each morning. In each condition data is from six separate mice. Orange and red dotted lines on the y-axis indicate the threshold for 10% and 20% (maximum limit) drops in body weight, respectively.

## SUPPLEMENTARY TABLES

**Supplementary Table. 1:** RP UPLC gradient elution in positive mode.

| Time (min) | Flow rate ($\mu\text{L}/\text{min}$) | Mobile Phase A (%) | Mobile Phase B (%) |
| --- | --- | --- | --- |
| 0.00 | 600 | 99.5 | 0.5 |
| 2.00 | 600 | 99.5 | 0.5 |
| 5.00 | 600 | 85.0 | 15.0 |
| 10.00 | 600 | 0.5 | 99.5 |
| 13.00 | 600 | 0.5 | 99.5 |
| 13.10 | 600 | 99.5 | 0.5 |
| 15.00 | 600 | 99.5 | 0.5 |

**Supplementary Table. 2:** MRM parameters for the RP UPLC-MS/MS NMN/NAD^+^ assay.

| Metabolite | Q1 | Q3 | RT (min) | DP | CE |
| --- | --- | --- | --- | --- | --- |
| NAD_1 | 664.0 | 136.0 | 0.88 | 70 | 70 |
| NMN_1 | 335.0 | 123.0 | 0.68 | 100 | 20 |

*Only main parent/daughter ions used in experiment are shown.*

**Supplementary Table. 3:** HILIC UPLC gradient elution in in positive mode.

| Time (min) | Flow rate (μL/min) | Mobile Phase A (%) | Mobile Phase B (%) |
| --- | --- | --- | --- |
| 0.00 | 500 | 95.0 | 5.0 |
| 1.00 | 500 | 95.0 | 5.0 |
| 8.00 | 500 | 50.0 | 50.0 |
| 9.00 | 500 | 50.0 | 50.0 |
| 9.10 | 500 | 95.0 | 5.0 |
| 15.00 | 500 | 95.0 | 5.0 |

**Supplementary Table. 4:**
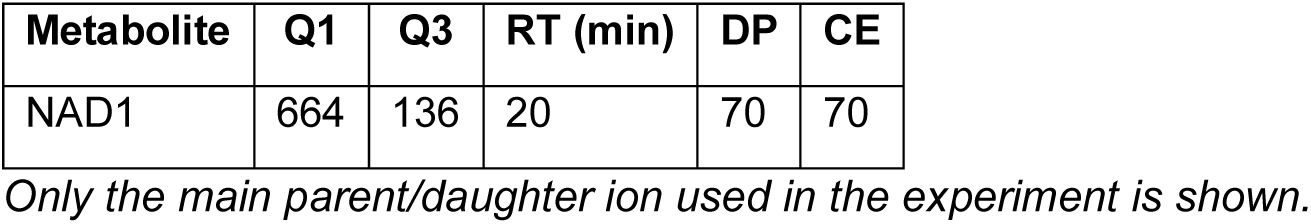
MRM parameters for the HILIC-positive UPLC-MS/MS NAD^+^ assay.

